# Programming the lymph node immune response with Amphiphile-CpG induces potent cellular and humoral immunity following COVID-19 subunit vaccination in mice and non-human primates

**DOI:** 10.1101/2022.05.19.492649

**Authors:** Lochana M. Seenappa, Aniela Jakubowski, Martin P. Steinbuck, Erica Palmer, Christopher M. Haqq, Crystal Carter, Jane Fontenot, Francois Villinger, Lisa K. McNeil, Peter C. DeMuth

## Abstract

Despite the success of currently authorized vaccines for the reduction of severe COVID-19 disease risk, rapidly emerging viral variants continue to drive pandemic waves of infection, resulting in numerous global public health challenges. Progress will depend on future advances in prophylactic vaccine activity, including advancement of candidates capable of generating more potent induction of cross-reactive T cells and durable cross-reactive antibody responses. Here we evaluated an Amphiphile (AMP) adjuvant, AMP-CpG, admixed with SARS-CoV-2 Spike receptor binding domain (RBD) immunogen, as a lymph node-targeted protein subunit vaccine (ELI-005) in mice and non-human primates (NHPs). AMP-mediated targeting of CpG DNA to draining lymph nodes resulted in comprehensive local immune activation characterized by extensive transcriptional reprogramming, inflammatory proteomic milieu, and activation of innate immune cells as key orchestrators of antigen-directed adaptive immunity. Prime-boost immunization with AMP-CpG in mice induced potent and durable T cell responses in multiple anatomical sites critical for prophylactic efficacy and prevention of severe disease. Long-lived memory responses were rapidly expanded upon re-exposure to antigen. In parallel, RBD-specific antibodies were long-lived, and exhibited cross-reactive recognition of variant RBD. AMP-CpG-adjuvanted prime-boost immunization in NHPs was safe and well tolerated, while promoting multi-cytokine-producing circulating T cell responses cross-reactive across variants of concern (VOC). Expansion of RBD-specific germinal center (GC) B cells in lymph nodes correlated to rapid seroconversion with variant-specific neutralizing antibody responses exceeding those measured in convalescent human plasma. These results demonstrate the promise of lymph-node adjuvant-targeting to coordinate innate immunity and generate robust adaptive responses critical for vaccine efficacy.

## INTRODUCTION

The evolving pandemic of coronavirus disease 2019 (COVID-19) has resulted in continued global health challenges with worldwide waves of infection driving significant morbidity and mortality despite widespread use of effective vaccines^1^. The serial emergence of viral variants of concern (VOC) with increased viral infectivity^2–4^, and mutations providing partial escape from neutralizing antibodies^5–8^ has proven a substantial challenge to successful control of viral spread. While currently authorized vaccines continue to provide valuable protection against hospitalization and severe disease, high community viral transmission has been observed despite high vaccination coverage^9,10^ Notably, prevention of severe disease appears to persist despite evidence of potentially rapid decay in vaccine-induced neutralizing antibody titers, and observations of reduced activity against recent VOC^9,11–14^. Therefore, while neutralizing antibody titers are considered the major correlate of protection for authorized vaccines,^15,16^ effective long-term protection against the worst outcomes of disease may be dependent on robust memory B and cross-reactive T cell responses capable of rapid re-activation to contain nascent viral infection in the absence of sterilizing immunity. Vaccines promoting responses with these immunological features may provide improved patient outcomes while also preventing the emergence of future coronavirus pathogens resulting from increasing exposure to natural zoonotic reservoirs.

We previously reported the development of ELI-005, a protein subunit vaccine combining the Spike receptor-binding-domain (RBD) protein with a lymph node targeting TLR-9 agonist, AMP-CpG, consisting of a diacyl lipid conjugated to CpG DNA^17^. The RBD immunogen contains targets of humoral and cellular immunity^18^ with a molecular size (34 kDa) predictive of lymph node accumulation following subcutaneous administration^17,19^. While conventional vaccine adjuvants of low molecular weight (<20 kDa) are readily absorbed into circulating blood following peripheral administration, AMP-modification is known to promote lymph node accumulation through non-covalent association with tissue albumin at the injection site^20,21^. AMP-directed biodistribution of vaccine components to draining lymph nodes can therefore significantly improve delivery to critical immune cells and engage mechanisms coordinating the magnitude and quality of the immune response while restricting exposure to immunologically irrelevant or tolerizing sites preferentially accessed by agents cleared through the blood. Previously, application of this approach in ELI-005 promoted highly potent RBD-specific T cell and neutralizing antibody responses following a three-dose immunization regimen in mice demonstrating the potential of lymph node targeted vaccination to generate immunity with an attractive balance of cellular and humoral responses^17^.

Here we describe further evaluation of ELI-005 in mice including the development of a simplified prime-boost regimen to rapidly induce VOC-directed cross-reactive antibody titers alongside highly potent T cell responses. Through comparison of soluble and AMP-CpG, we explore the differential underlying mechanisms of innate immune response activation in draining lymph nodes and their dependence on effective lymph node targeting of adjuvant. We also evaluated ELI-005 for T cell and humoral immunogenicity in Rhesus macaques as a predictive model for potential safety and activity in human subjects.

## RESULTS

### AMP-CpG promotes strong and durable cellular and humoral anti-SARS-CoV-2 responses in mice

Prior evaluation of ELI-005 demonstrated potent induction of RBD-specific immunity characterized by high frequency peripheral and tissue resident T cell responses alongside durable neutralizing antibody titers following a three-dose immunization regimen in mice^17^. To improve the feasibility of ELI-005 immunization for potential clinical translation, we evaluated a prime-boost regimen administered on weeks 0 and 2, with 10 µg Wuhan-Hu-1 (WH-01) SARS-CoV-2 Spike RBD admixed with CpG DNA adjuvant (Figure 1a). To assess the importance of AMP-modification on the magnitude and longevity of induced immune responses, RBD antigen was administered with 1 nmol of AMP-CpG (consisting of diacyl lipid conjugated to CpG-7909 DNA) or the unmodified soluble CpG analog. Cellular and humoral responses were assayed one week after the boost dose to characterize acute responses, and longitudinally over the course of 32 weeks to assess long-term immune persistence as depicted in Figure 1a. Mice immunized with ELI-005, including AMP-CpG, had a 12-fold increase in the number of splenic IFNγ spot-forming cells (SFCs) compared to mice treated with soluble CpG (Figure 1b). Similarly, compared to vaccination with soluble CpG, AMP-CpG induced significantly increased RBD-specific cytokine-producing CD8^+^ T cells in peripheral blood following stimulation with overlapping RBD peptides directly ex vivo (Figure 1c). Assessment of CD4^+^ T cells in peripheral blood showed a similar hierarchy of response with only AMP-CpG immunization eliciting significantly increased frequencies of RBD-specific cytokine^+^ cells (Figure 1d). Similar trends were observed in CD8^+^ and CD4^+^ T cells isolated from perfused lung tissue, an important site for resident T cell immunity to facilitate early detection and mediation of nascent viral infection (Figure 1e-f). Given the strength of AMP-CpG to promote significantly enhanced peak immune responses following a prime-boost regimen, and the importance of long-lived immune-protection, longitudinal measurement of immunogenicity was performed for up to 32 weeks. Mice immunized with AMP-CpG showed higher levels of circulating RBD-specific CD8^+^ cytokine^+^ T cells throughout the assessment period, compared to animals immunized with soluble CpG or mock (Figure 1g-h). Together, these results demonstrate the potency of ELI-005 prime-boost immunization and the critical role of AMP-CpG to promote significantly enhanced and persistent cellular immunity in multiple tissues important for potential prophylactic protection or moderation of disease severity.

**Fig. 1.**
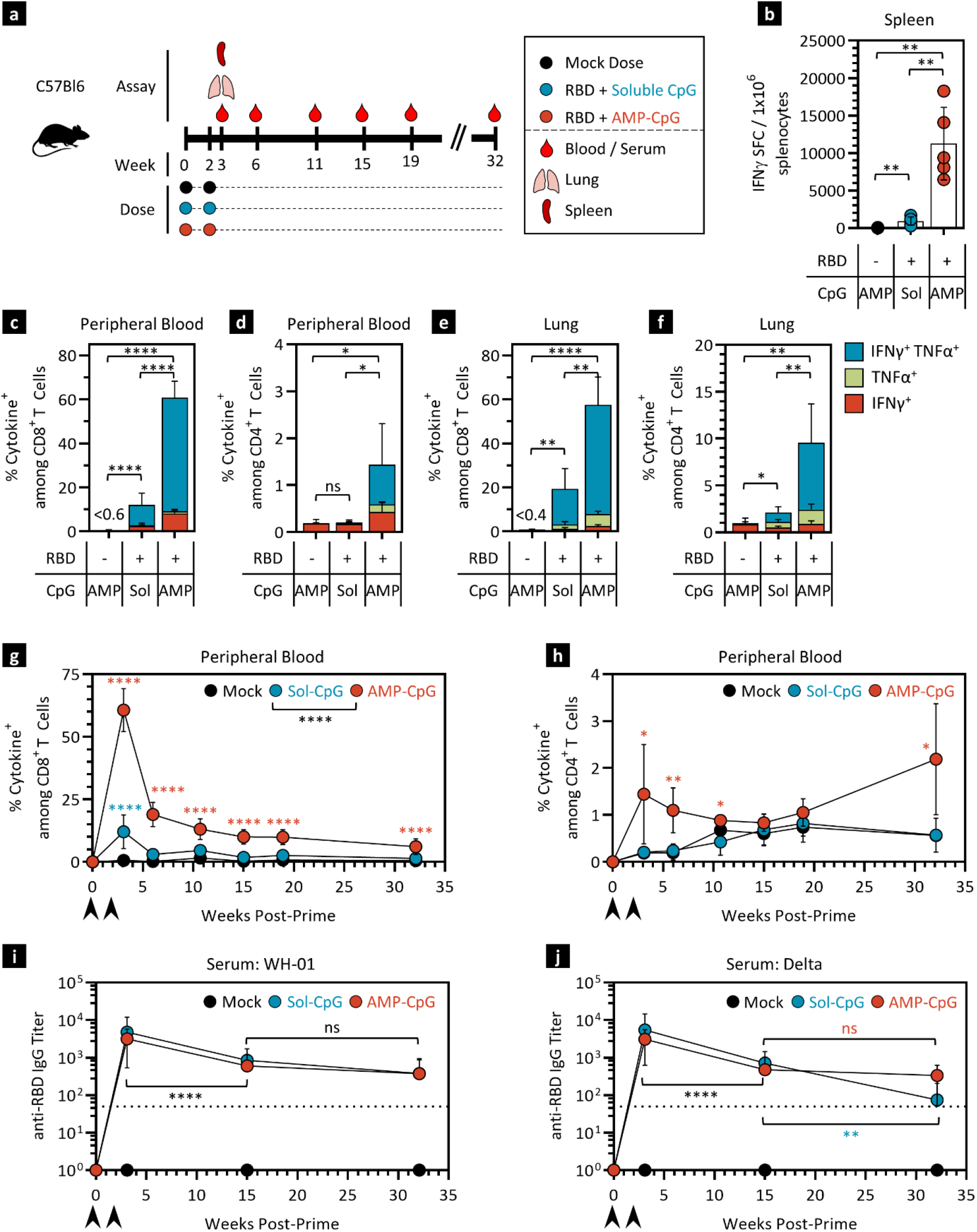
AMP-CpG immunization elicits potent, long-lived immunity to SARS-CoV-2 in mice. **a** Schema showing animal dosing and experimental schedule. C57Bl/6J mice were immunized twice with 10 µg WH-01 Spike RBD protein and 1 nmol soluble or AMP-CpG. **b** Splenocytes were restimulated with Spike RBD OLPs 7 days post booster dose and assayed for IFNγ production by ELISpot. Shown is frequency of IFNγ SFCs per 1 × 10^6^ splenocytes. **c-h** Flow cytometric analysis of cytokine production by CD8^+^ (**c, e, g**) and CD4^+^ T cells (**d, f, h**). Shown are percentages of cytokine^+^ cells among CD8^+^ or CD4^+^ T cells: peak responses 7 days post booster dose in peripheral blood (**c-d**), and perfused lung tissue (**e-f**), and longitudinal analysis over 32 weeks in peripheral blood (**g-h**). **i-j** Serum anti-RBD IgG titers were determined against WH-01 RBD (**i**) and Delta RBD (**j**) protein. Values depicted are means ± standard deviations. \**p* < 0.05, \*\**p* < 0.01, \*\*\**p* < 0.001, \*\*\*\**p* < 0.0001 by two-sided Mann-Whitney or t-test applied to cytokine^+^ T cell frequencies or antibody titers.

Long-term evaluation of the humoral response showed rapid seroconversion after prime-boost immunization for both soluble and AMP-CpG immunized groups. Serum IgG antibody titers against both the immunizing antigen WH-01 RBD, as well as the Delta RBD, peaked shortly after boost immunization and were maintained at high levels for the duration of this study (Figure 1i-j). Overall, these data indicate that ELI-005, adjuvanted with AMP-CpG, generates potent cross-reactive humoral immune responses that persist long term at significant levels.

### AMP-CpG induces T cell memory responses which rapidly expand upon antigen re-exposure

The persistence of circulating RBD-specific T cells post-boost suggests that ELI-005 promotes the development of durable memory T cells. Analysis of this long-lived T cell population showed a significantly larger population of circulating RBD-specific tetramer^+^ cells with a CD44^+^CD62L^+^ central memory phenotype among CD8^+^ T cells. AMP-CpG vaccine-induced responses were 2.5-fold greater than in the soluble CpG comparator group (Figure 2a), suggesting the potential for a more rapid expansion of effector T cells upon future exposure to SARS-CoV-2. Consequently, we characterized the recall response upon antigen re-exposure following subcutaneous administration of 10 µg WH-01 RBD, 17 weeks after boost. Recall responses in circulating blood analyzed 7 days later confirmed rapid expansion of RBD-specific CD8^+^ T cells in ELI-005 immunized animals (14-fold from 1.9% pre-challenge to 26.8% post recall, Figure 2b). In contrast, recall responses in soluble CpG immunized mice were significantly lower, indicating a substantial advantage for long-term memory response expansion in AMP-CpG immunized animals.

**Fig. 2.**
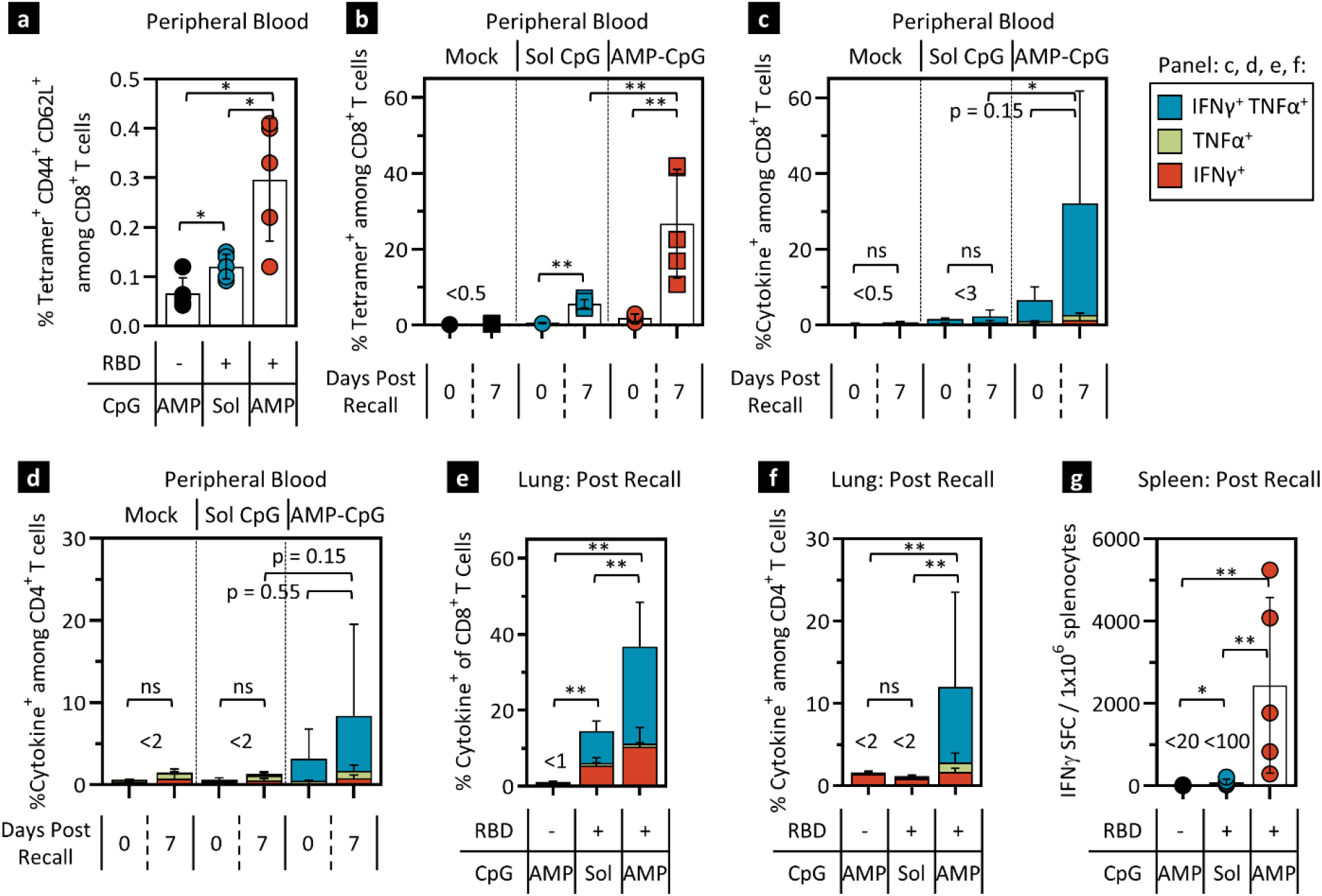
AMP-CpG immunization generates long-lived memory T cell responses that rapidly expand upon antigen re-exposure in mice. C57Bl/6J mice (n=5) were immunized twice with 10 µg WH-01 RBD protein and 1 nmol soluble or AMP-CpG. **a-b** 17 weeks post dose 2, blood was collected for a pre-recall time point, followed by subcutaneous challenge with 10 µg WH-01 RBD protein on the next day, and a second blood collection 7 days later. **a** Pre-challenge peripheral blood leukocytes were stained for CD44 and CD62L. Shown are percentages of tetramer^+^ CD44^+^ CD62L^+^ cells among CD8^+^ T cells. **b** Peripheral blood CD8^+^ T cells collected before and after challenge were stained with tetramer specific for Spike RBD and analyzed by flow cytometry. **c-g** 30 weeks post dose 2, mice were challenged subcutaneously with 10 µg WH-01 RBD protein and assayed 7 days later. CD8^+^ T cells collected from blood (**c**) and perfused lung (**e**), and CD4^+^ T cells collected from blood (**d**) and perfused lung (**f**) were stimulated with WH-01 RBD OLPs and assayed for intracellular cytokines by flow cytometry. **g** Splenocytes were restimulated with WH-01 RBD OLPs and assayed for IFNγ production by ELISpot. Mock vaccines contained AMP-CpG without the addition of antigen. Values depicted are means ± standard deviation. *\*p < 0*.*05, **p < 0*.*01* by two-sided Mann-Whitney or t-test applied to tetramer^+^ or cytokine^+^ T cell frequencies, or SFC numbers.

Considering the robust T cell recall response observed 17 weeks after boost, an additional cohort of animals was recalled 30 weeks after boost. Circulating CD8^+^ and CD4^+^ T cell responses were measured prior to and 7 days post recall with 10 µg WH-01 RBD antigen. In the peripheral blood of animals immunized with AMP-CpG, the percentage of polyfunctional cytokine-producing CD8^+^ and CD4^+^ T cells increased 5-fold to 32% and nearly 3-fold to 8.5% post challenge, respectively (Figure 2c-d). This compared to just 2.3% of CD8^+^ and 1.3% of CD4^+^ T cells that displayed a cytokine^+^ phenotype in the soluble CpG group. Of note, the long-term CD8^+^ and CD4^+^ responses 32 weeks after immunization with AMP-CpG persisted at levels that were higher than responses in the soluble CpG group even after they had been recalled.

Cellular immune responses were analyzed in tissues important for point-of-entry and systemic protection against SARS-CoV-2. Perfused lung tissue and spleens from immunized mice were analyzed 7 days after WH-01 RBD challenge at week 32. Antigen recall of animals originally immunized with AMP-CpG induced a large proportion of lung resident T cells to secrete polyfunctional cytokines. In these animals, 37% of CD8^+^ T cells (Figure 2e) and 12% of CD4^+^ T cells (Figure 2f) secreted cytokines, compared to 14% and <2% in mice treated with soluble CpG, respectively. Similar trends were observed in spleen, where immunization with AMP-CpG produced a mean frequency of nearly 2,500 IFNγ SFC per 10^6^ splenocytes opposed to < 100 IFNγ SFC per 10^6^ splenocytes in the soluble CpG group. These data indicate that AMP-CpG-adjuvanted ELI-005 generates robust and durable T cell memory populations that can quickly and potently expand upon re-exposure to SARS-CoV-2 antigens, providing potent, long-term cellular immunity at important sites for immune surveillance and disease moderation. Humoral responses did not show an increase in serum titer at 7 days post recall, suggesting that the antibody recall response may lag behind that of T cells (Supplemental Figure S1a-b).

### ELI-005 is highly stable for extended periods of time

The previous data demonstrated the potency and durability of ELI-005-induced immunity. However, global vaccine distribution in support of mass immunization campaigns requires stability at near ambient temperature without dependence on an ultra-cold storage supply chain. To examine the stability of ELI-005, an immunogenicity study was conducted, in which formulated vaccine doses were stored at common refrigerator temperatures (4°C) for 2-4 weeks or at room temperature (22°C) for 3 days prior to dosing. One week after the second immunization, animals were assessed for their cellular and humoral immune responses. Cytokine producing CD8^+^ and CD4^+^ T cells in both peripheral blood and lung, as well as serum antibody titers, demonstrated near-identical trends to those observed in Figure 1 where doses were prepared freshly before injection (Supplemental Figure S2a-e). This suggests the potential for ELI-005 to maintain immunogenicity at storage conditions that would enable effective global distribution without costly and impractical logistical demands.

### AMP-CpG stimulates comprehensive transcriptional programming of immune orchestration in draining lymph nodes

The enhanced immunogenicity of ELI-005 is correlated to past observations of improved AMP-CpG delivery to draining lymph nodes *in vivo*^20,21^. To further explore the resulting immune biology, a comprehensive panel of 568 immune-relevant gene transcripts was assessed in draining lymph nodes using Nanostring. Mice were immunized once with WH-01 RBD admixed with soluble or AMP-CpG and draining inguinal lymph nodes were assessed for differential regulation of gene transcription at the indicated time points (Figure 3a). Mice treated with AMP-CpG exhibited an acute and significant upregulation of inflammatory chemokine mRNA as early as 2 hours post-immunization (Supplemental Figure S3a-c), including *Ccl3* (MIP-1α), *Ccl12* (MCP-5) and *Cxcl10* (IP-10), important chemo-attractants for monocytes, macrophages, and dendritic cells (DCs). This trend was continued at 6 hours, when additional chemokines *Ccl7* (MCP-3) and *Ccl9* (MIP-1γ) were detected, potentially enhancing the chemotactic recruitment of professional APCs to the lymph nodes (Figure 3b and Supplemental Figure S3d-e). In contrast, mice treated with soluble CpG displayed a very limited chemokine transcriptional profile restricted to only *Ccl12* (MCP-5). AMP-CpG treated lymph nodes exhibited comprehensive transcriptional activation at 24 hours, reflecting multiple axes of robust innate immune activity critical to downstream adaptive immunogenicity (Figure 3c-e). Of specific interest were transcriptional signatures suggesting an influx of APCs, poised for pathogen recognition (e.g., *Tlr3/4/9, Myd88*, cytosolic PRRs), antigen processing and presentation (e.g., cathepsins, *Tap1, β2m*), and co-stimulation (eg., *Cd40, Cd86*). Transcriptional evidence of innate immune cell recruitment and activation in the lymph nodes was further associated with upregulated inflammatory (e.g., *Tnf, Stat3*), and anti-viral (e.g., *Stat1*, IRFs) processes suggesting a potential role for these pathways in AMP-CpG-driven acute immune activation. In contrast, minimal transcriptional changes were induced in the lymph nodes by soluble CpG consistent with prior evidence showing poor lymph node accumulation and reduced immunogenicity *in vivo*. Assessment at 72 hours post immunization indicated a peak in soluble CpG-mediated transcriptional changes dominated by indicators of B cell recruitment and activation (e.g., *Cd19, Cd79a, Btk*, Figure 3f, Supplemental Figure S3f-g). In contrast, AMP-CpG immunized lymph nodes showed minimal B cell associated transcriptional patterns at this time point. Instead, analysis showed evidence of subsiding pattern recognition- and chemokine-encoding transcript levels, but persistent signatures of upregulated antigen presentation and inflammatory pathways consistent with lymph node mobilization of professional APCs to coordinate downstream adaptive immunity. Overall, these data indicate a striking contrast between soluble- and AMP-CpG-induced transcriptional programming. This demonstrates the critical importance of AMP-modification to affect the conversion of the naïve lymph node into a highly inflammatory and anti-viral niche, enabling APCs to detect, process, and present antigens to the adaptive immune system, culminating in robust downstream adaptive immunity.

**Fig. 3.**
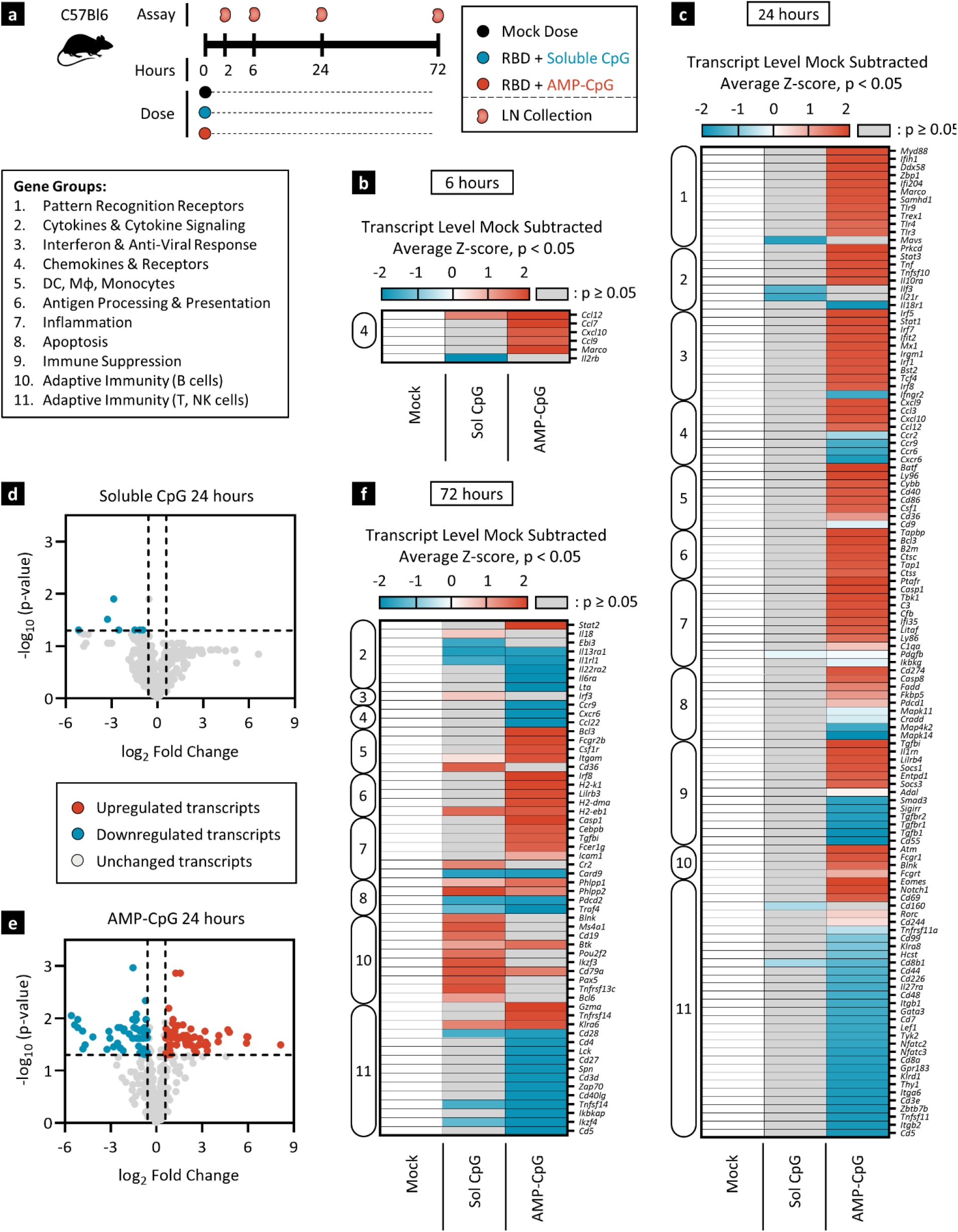
AMP-CpG immunization induces lymph node transcriptional reprogramming reflecting APC recruitment with increased potential for antigen processing and presentation. C57Bl/6J mice (n=2) were immunized once with 10 µg WH-01 RBD protein and 1 nmol soluble or AMP-CpG. **a** Schema showing animal dosing and experimental schedule. **b, c, f** Heatmap representation of whole lymph node mRNA analyzed by NanoString nCounter® Mouse Immunology Panel. Shown are mock-subtracted, average Z-scores of gene transcript levels significantly (p < 0.05) downregulated (≥ -1.5-fold change, blue) or upregulated (≥ 1.5-fold change, red) at 6 hours (**b**), 24 hours (**c**), and 72 hours (**f**) post injection relative to mock immunization. Insignificant values with p ≥ 0.05 are shown in gray. Genes were clustered into 11 groups (box insert) using Gene Ontology databases. **d-e** Volcano plot representation of log-transformed transcript values from soluble CpG (**d**) and AMP-CpG (**e**) immunized animals at 24 hours depicted in **c**. Mock vaccines contained PBS vehicle only. Statistical analysis was performed using Rosalind software. Dotted horizontal line represents significance threshold of p = 0.05; vertical dotted lines represent fold-change limits of ± 1.5-fold change. LN: lymph node.

### AMP-CpG induces potent acute activation of lymph node proteomic and cellular innate immune responses

Given the evidence of AMP-CpG-mediated transcriptional reprogramming in the lymph nodes, we further characterized the lymph node proteomic milieu and immune cell composition to further investigate the critical mechanisms underlying the induction of adaptive immunity following vaccination. For this purpose, we utilized two approaches: first, the total expression of inflammatory proteins induced by immunization with soluble or AMP-CpG was measured in lymph node lysates by multiplexed proteomics; second, lymph node immune cell numbers, activation markers and intracellular cytokines were assessed by flow cytometry. Draining lymph nodes from mice immunized with either soluble or AMP-CpG were collected 6 and 24 hours after prime immunization and lysates were assessed by Luminex in comparison to animals receiving mock injections of vehicle only (Figure 4a-b). At 6 hours soluble and AMP-CpG immunizations induced qualitatively similar increases in several protein analytes, including chemokines associated with innate cell recruitment and spatial lymph node organization of innate-adaptive immune cell interactions (IP-10, MCP-1, MIP-1α, MIP-1β; Figure 4a, Supplemental Figure S4a). Assessment at 24 hours post-vaccination revealed that while responses to soluble CpG immunization had almost universally reverted to baseline levels, those induced by AMP-CpG persisted and expanded to include numerous factors associated with innate-cell mediated orchestration of adaptive immunity (Figure 4b, Supplemental Figure S4b). AMP-CpG immunization resulted in robust and comprehensive elevation of nearly all analytes tested, including growth factors, Th1-associated pro-inflammatory cytokines, Th-2 and regulatory cytokines, chemokines, inflammasome-associated cytokines, and Type I interferon. Given the observation that AMP-CpG induces a strong lymph node pro-inflammatory milieu, we further assessed the importance of contributions from various immune cells to orchestrate this process in the lymph nodes 6, 24, and 48 hours following prime immunization (Figure 4c-g, line graphs, Supplemental Figure S4c-g). Analysis by flow cytometry at 24 hours (Supplemental Figure 5) showed that AMP-CpG, but not soluble CpG, significantly increased the total numbers of all analyzed innate cell lineages, including monocytes, macrophages, neutrophils, DCs, and NK cells. Notably, monocyte and neutrophil cell numbers peaked at 24 hours in AMP-CpG immunized animals and decreased thereafter, consistent with their role in acute responses, while macrophages, DCs and NK cell numbers continued to increase at 48 hours. Importantly, while soluble CpG immunization resulted in similar elevations in total lymph node populations of macrophages, DCs, and NK cells, AMP-CpG universally promoted a significantly stronger profile of innate cell activation status as indicated by the number of lymph node resident cells producing multiple key cytokines (IFNγ, IFNβ, IL-1β, IL-6, IL-12) and elevated CD86 surface expression at 24 hours (Figure 4c-g, heat maps, Supplemental Figure S4c-g). Together with the observed lymph node proteomics profile, these data indicate the essential role of AMP-modification to enable lymph node delivery and potent innate immune activation by CpG DNA *in vivo*.

**Fig. 4.**
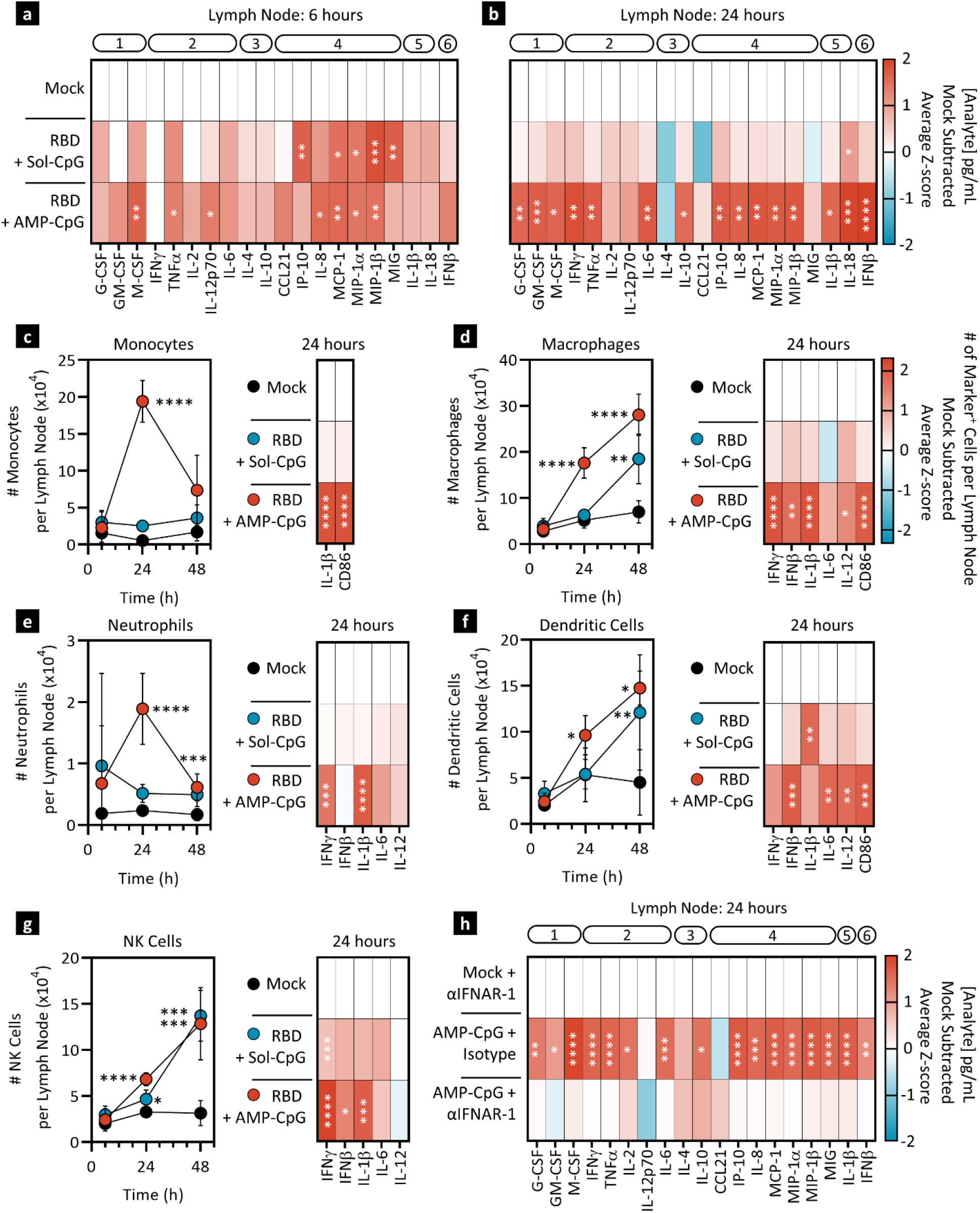
AMP-CpG immunization induces lymph node proteomic signatures of immune activation with multiple innate cell lineage involvement. **a-g** C57Bl/6J mice (n=5) were immunized with 10 µg WH-01 Spike RBD protein and 1 nmol soluble or AMP-CpG. Mock immunized animals received PBS vehicle only. Lymph nodes were collected and processed for protein extraction (**a-b**) or flow cytometry (**c-g**). Lymph nodes collected at 6 hours (**a**) and 24 hours (**b**) post immunization were homogenized and extracted protein levels were measured by Luminex. Results are expressed as heat maps of mock subtracted Z-scores of analyte concentrations (pg/ml). Analytes are annotated by functional category: (1) growth factors, (2) Th1 cytokines, (3) Th2/regulatory cytokines, (4) chemokines, (5) inflammasome, and (6) Type I interferon. **c-g** Lymph nodes were collected at 6, 24 and 48 hours and single cell suspensions were analyzed by flow cytometry. Line graphs represent cell subset numbers per lymph node across time. Heat maps are mock-subtracted Z-scores of specific cell subsets expressing the respective marker per lymph node. **h** C57Bl/6J mice (n=5) were dosed intraperitoneally with IFNAR-1 blocking antibody one day prior to immunization with AMP-CpG vaccine. Mock immunized animals received PBS vehicle only. Protein from lymph nodes was extracted 24 hours later. Luminex assessed analyte concentrations were expressed in heat maps showing mock-subtracted Z-scores of analyte concentration (pg/ml) and annotated as in **a-b**. Values depicted are mean ± standard deviation. *\*p < 0*.*05; **p < 0*.*001; ***p < 0*.*0005; ****p < 0*.*0001* by one-way ANOVA comparison of experimental groups to mock control.

**Fig. 5.**
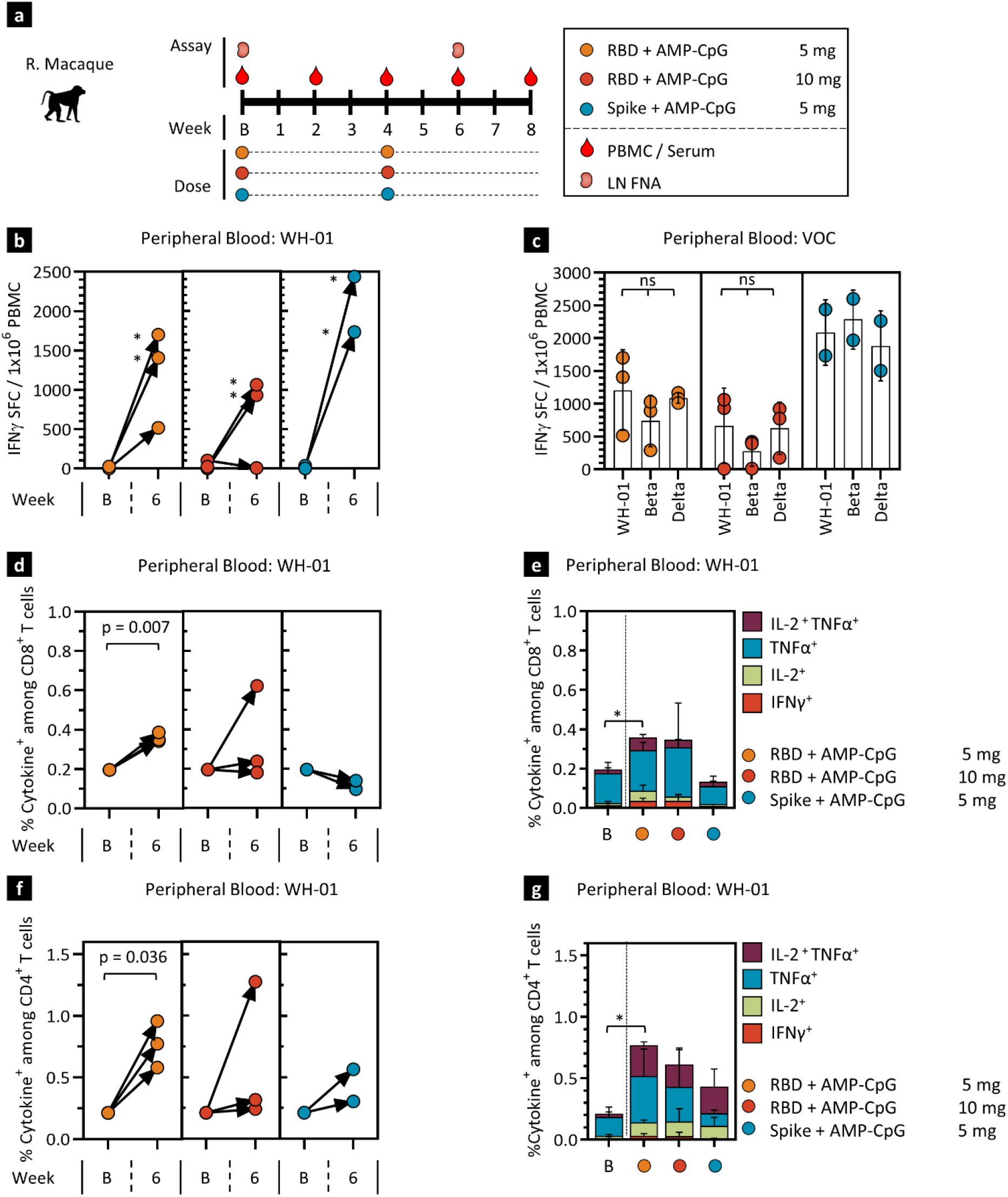
AMP-CpG immunization induces robust T cell responses to multiple variants of SARS-CoV-2 in NHPs. Rhesus macaques (n=2/3) were immunized at week 0 and 4 with 140 µg WH-01 RBD protein admixed with either 5 mg or 10 mg of AMP-CpG, or 140 µg full WH-01 Spike protein admixed with AMP-CpG at 5 mg. PBMCs were collected at baseline and week 6 for T cell response assessment. **a** Schema showing animal dosing and experimental schedule. **b-c** PBMCs were stimulated at 0 and 6 weeks with WH-01 RBD OLPs (**b**) or with VOC OLPs (**c**) at 6 weeks prior to analysis by IFNγ ELISpot assay. Shown are IFNγ SFCs per 1 × 10^6^ PBMCs. **d-g** PBMCs were stimulated with WH-01 OLPs prior to analysis for intracellular cytokines by flow cytometry. Shown are frequencies of total (**d, f**) and individual IL-2, TNFα, IFNγ cytokine producing (**e, g**) CD8^+^ (**d, e**) and CD4^+^ T cells (**f, g**). Values depicted are mean ± standard deviation. *\* p < 0*.*05; ** p < 0*.*01* by two-sided Mann-Whitney test applied to cytokine^+^ T cell frequencies. LN: lymph node, B: baseline

Upon observing the induction of IFNβ, the strong transcriptional signature of interferon response and apparent involvement of all major innate immune cell lineages in the lymph node immune response following immunization with AMP-CpG, we hypothesized that activation might proceed in part through stimulation of the Type I interferon antiviral response pathway. To investigate the role of this signal cascade in the innate responses induced by AMP-CpG, mice were pre-treated with an IFNAR-1 blocking mAb to abrogate both IFNα and IFNβ signal transduction. Mice were then immunized, and lymph nodes collected after 24 hours for proteomics analysis. While control mice pretreated with inactive isotype control antibody displayed elevation of nearly all measured analytes, the AMP-CpG-induced inflammatory milieu in the lymph node was completely abrogated by IFNAR-1 blockade (Figure 4h, Supplemental Figure S4h), confirming a critical role for this pathway in AMP-CpG-induced immune activation.

### AMP-CpG stimulates broadly reactive cellular immune responses targeting multiple VOC in non-human primates (NHPs)

Motivated by the robust cellular and humoral immunity observed following ELI-005 immunization in mice, we conducted immunogenicity assessments in NHPs to evaluate activity predictive of potential responses in human subjects. Immunogenicity was assessed following a prime-boost regimen in 8 adult female rhesus macaques (Figure 5a), 4-5 years old, following immunization with 140 µg WH-01 RBD admixed with either 5 mg (n=3) or 10 mg (n=3) of AMP-CpG, or 140 µg of full WH-01 Spike protein admixed with 5 mg of AMP-CpG (n=2). Safety assessments showed no adverse events or signs of potential clinical risk as determined through longitudinal monitoring of body temperature, body weight, blood cell counts, serum cytokine measurement, and gross clinical observations including monitoring of the local injection site (Supplemental Figure S6 and data not shown). Throughout the study, peripheral blood mononuclear cells (PBMCs), serum, and lymph node fine-needle aspirates (FNA) were collected for analysis of cellular and humoral immune responses (Figure 5a). At week 6, two weeks following the boost immunization, individual animals from all vaccine groups exhibited significantly elevated frequencies of IFNγ SFC in PBMC after stimulation *ex vivo* with WH-01 RBD overlapping peptides (OLPs), with most animals having greater than 1,000 SFC per 10^6^ PBMC (Figure 5b). Potential cross-reactivity of the induced responses was evaluated through stimulation with VOC-specific OLPs, and no significant differences in the frequency of IFNγ SFC were observed for Beta and Delta OLP stimulation as compared to WH-01 (Figure 5c), suggesting that ELI-005-induced T cell responses may target epitopes conserved across VOC sequences. Additional assessment of PBMC-derived T cell responses was completed through analysis of multi-cytokine production after stimulation *ex vivo*. Among immunization groups, WH-01 RBD admixed with 5 mg of AMP-CpG elicited the most robust peripheral blood T cell responses at week 6 with 0.45% of CD8^+^ T cells (Figure 5d-e) and 0.8% of CD4^+^ T cells (Figure 5f-g) observed to produce IL-2, TNFα, IFNγ, or combinations of these upon stimulation. Other vaccine cohorts showed relatively lower cytokine^+^ CD8^+^ T cell responses but cytokine^+^ CD4^+^ responses trended consistently higher suggesting that CD4^+^ T cell responses are more dominant in NHPs than previously observed for mice.

### AMP-CpG elicits lymph node germinal center (GC) development and rapid induction of cross-reactive neutralizing antibody responses

At week 6, all vaccine combinations induced RBD-specific GC B cells (RBD^+^ CD20^+^ BcL6^+^ Ki-67^+^) with elevated frequencies compared to baseline levels in the lymph nodes (Figure 6a-b). Serum was assessed for RBD-specific IgG at longitudinal timepoints throughout the study. Evaluation of serum IgG specific for WH-01 RBD showed rapid and nearly universal seroconversion across all RBD-immunized treatment groups as early as week 2 and 4 after the initial immunization (Figure 6c). Further increases in IgG serum concentrations were observed for all RBD-immunized groups post boost at weeks 6 and 8 with peak responses at week 6 elevated up to 5,000-fold relative to baseline. Similar trends were observed for IgG responses following immunization with full Spike protein suggesting that both RBD and full-length Spike are immunogenic in combination with AMP-CpG adjuvant.

**Fig. 6.**
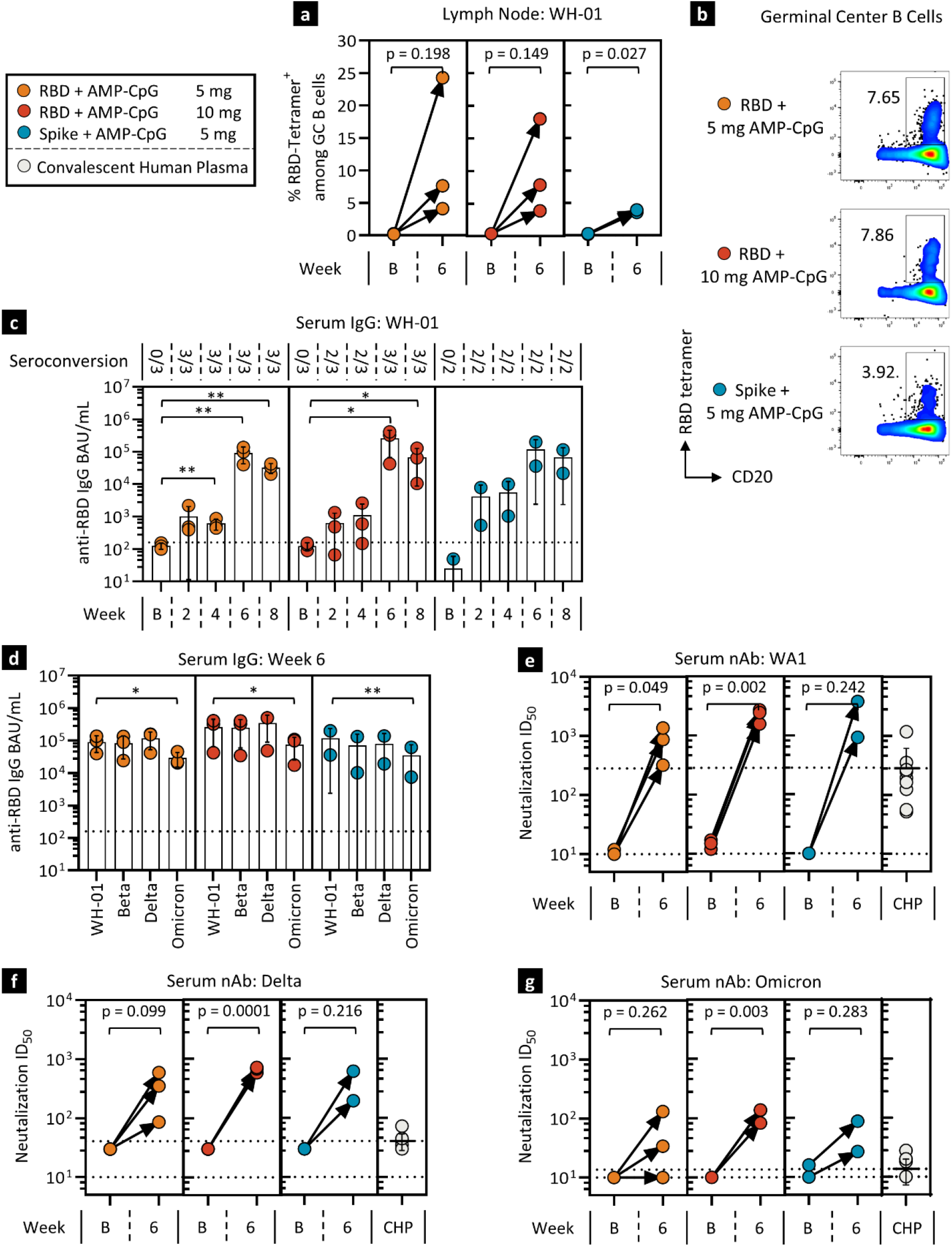
AMP-CpG immunization stimulates rapid and potent humoral responses to multiple variants of SARS-CoV-2 in NHPs. Rhesus macaques were immunized as shown in **fig 5a** and humoral responses to SARS-CoV-2 were assessed in lymph node FNA and serum at various timepoints. **a-b** Shown are frequencies of RBD-specific GC B cells (CD20^+^ BcL6^+^ Ki-67^+^) in lymph node FNAs at baseline and week 6 with corresponding representative flow cytometry dot plots of RBD-specific B cells. **c-d** RBD-specific serum IgG binding unit concentrations were assessed for WH-01 (**c**) and VOC (**d**) RBD-binding activity, either longitudinally or at baseline and week 6. **e-g** Neutralizing antibody responses were determined through pseudovirus inhibitory activity. Shown are ID_50_ values for WA1 (**e**), Delta (**f**), and Omicron (**g**) in comparison to convalescent human plasma (CHP). Values depicted are means ± standard deviation. The dotted lines represent the lower limit of detection discriminating between the samples positive or negative for seroconversion and the mean value observed for the human plasma comparators where appropriate. \**p* < 0.05; \*\**p* < 0.01 by two-sided Mann-Whitney test.

Antibody cross-reactivity was assessed to evaluate the potential for induced IgG responses to recognize a panel of RBD variants associated with VOC driving recent pandemic waves of infection. Immunization with AMP-CpG elicited serum IgG responses with broad cross-reactivity targeting several predominant VOC. At week 6, across all RBD-immunized groups, no significant decrease in RBD-specific IgG serum concentration was observed for Beta or Delta RBDs relative to WH-01 (Figure 6d). IgG reactivity to Omicron RBD was robust though decreased relative to Wuhan, Beta and Delta. Similar trends were seen for animals immunized with full Spike immunogen.

Consistent increases in the level of virus-specific neutralizing antibodies against WH-01 were observed in serum from all immunized animals at week 6 with neutralization ID_50_ approximately 10-fold higher than levels observed in a panel of convalescent human plasma samples (Figure 6e). Week 6 neutralizing antibody responses against Delta and Omicron variants were also consistently elevated relative to baseline but reduced in comparison to WH-01-specific responses targeting the vaccine immunogen (Figure 6f-g). However, these variant-directed responses, including those specific to Omicron, consistently trended higher than the convalescent human plasma comparators suggesting AMP-CpG vaccine-induced cross-reactive neutralizing responses may exceed levels commonly induced by natural infection in humans.

### AMP-CpG induces robust pro-inflammatory transcriptional programming in draining lymph nodes of NHPs

To survey signaling patterns in AMP-CpG vaccinated NHPs, lymph node FNA were collected at baseline and 24 hours post-immunization for transcriptional analysis of immune response using Nanostring. Dynamic changes in numerous gene transcript levels were observed across various gene ontology categories spanning multiple axes of immune coordination (Figure 7a). Overall, 44 genes were differentially expressed relative to baseline; of these 27 were significantly upregulated at least 1.5-fold after vaccination and 17 genes were downregulated at least 1.5-fold (Figure 7b). Upregulated transcripts included genes involved in adaptive and innate immunity, APC activation, interferon signaling and migration while genes involved in T cell anergy and exhaustion were highly downregulated. Notably, *Cd83* and *Batf3*, transcripts important for DC activation and cross-presentation of antigens to CD8^+^ T cells^22,23^, were highly induced as was the pro-inflammatory Th1 cytokine, *Il18* (Figure 7c). AMP vaccination showed strong induction of genes principally involved in interferon signaling including *Stat1* and *Ifnar2*. The largest category of upregulated genes was involved with migration, including *Cxcl10* (IP-10) which is chemotactic for macrophages, T cells, NK cells and DCs, and *Ccl3* (MIP-1α) which orchestrates T cell-APC encounters in the lymph nodes to enhance the magnitude of memory T cells^24–26^. In contrast, genes involved in T cell anergy and exhaustion were downregulated including *Siggir* which inhibits signal transduction of TLRs, *Il18r1* and *Il1r1* to negatively regulate inflammation^27^. These results share notable similarities to those collected in mice and further confirm that immunization in NHPs with AMP-CpG induces a pro-inflammatory environment to promote adaptive immunity including T cell priming and activation in the draining lymph nodes.

**Fig. 7.**
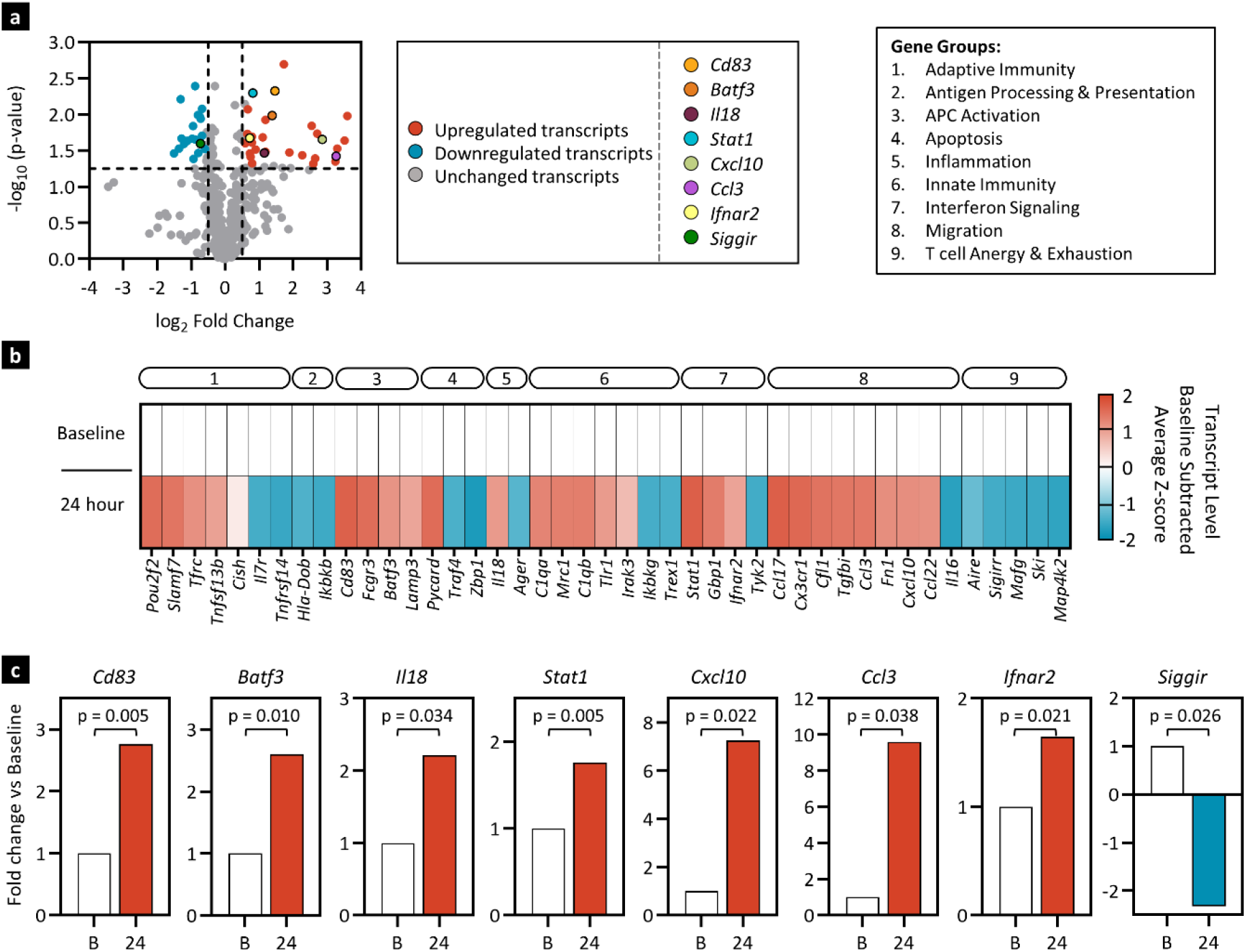
AMP-CpG immunization stimulates dynamic changes in immune gene transcript levels in NHP draining lymph nodes. Rhesus macaques (n=3) were immunized at week 0, 4 and 10 with 140 µg WH-01 RBD admixed with 5 mg of AMP-CpG. Lymph node FNAs were collected at baseline and 24 hours post-week 10 boost. Nanostring analysis, using the NanoString nCounter® NHP Immunology Panel, was performed comparing gene expression in lymph node FNA between baseline and 24 hours post boost. **a** Volcano plot representation of log-transformed transcript values of 770 genes at 24 hours. Dotted horizontal line represents significance threshold of p = 0.05; vertical dotted lines represent fold-change limits of ± 1.5-fold change. **b** Heat map of gene transcripts that were differentially expressed. Shown are mock-subtracted, average Z-scores of gene transcript levels significantly (p < 0.05) downregulated (≥ -1.5-fold change, blue) or upregulated (≥ 1.5-fold change, red) relative to baseline values. Insignificant values with p ≥ 0.05 are shown in gray. Genes were clustered into 9 groups (box insert) using Gene Ontology databases and annotated at the top of the heat map: (1) adaptive immunity, (2) antigen processing and presentation, (3) APC activation, (4) apoptosis, (5) inflammation, (6) innate immunity, (7) interferon signaling, (8) migration, and (9) T cell anergy and exhaustion. **c** Fold-change bar graphs for selected gene transcripts. Statistical analysis was performed using Rosalind software.

## DISCUSSION

Despite authorization and widespread uptake of effective vaccines, the COVID-19 pandemic continues to result in global waves of infection driven by the emergence of additional variants^28^, and relatively short-lived sterilizing immunity acquired through natural infection or immunization^29–31^. As viral variants progressively develop more substantial mutational deviation from ancestral strains and acquire greater infectivity, the objective for future vaccine candidate activity may shift from prevention of viral infection by neutralizing antibodies to T cell-mediated prevention of progression to severe disease, hospitalization, and death. While currently approved vaccines have shown rapidly diminishing neutralizing antibodies, T cell responses are potentially more durable^32,33^ while also providing potential for broader current and future VOC recognition^31,34,35^. Thus, vaccines capable of promoting balanced humoral and cellular immunity providing long-term persistence, broad cross-reactive viral recognition, and rapid memory re-activation upon recall will be attractive candidates for further development.

Accordingly, prime-boost immunization with ELI-005 in mice and NHPs induced parallel T cell and antibody immunity. Antibody responses in NHPs exceeded the ability of convalescent human plasma to neutralize multiple VOCs. Importantly, Omicron-specific neutralizing responses were detected after only two doses, suggesting an advantage over currently approved vaccines where a third dose has proven necessary to induce similar responses in humans^36,37^. Further, ELI-005-induced antibody responses in NHPs were accompanied by robust development of lymph node GC B cell responses thought to correlate with prolonged memory B cell responses in human vaccinees^38,39^. These responses, exhibiting potential for broadly cross-neutralizing humoral recognition and induction of GC precursors of durable memory B cell responses, may provide immunity with improved persistence and relevant specificity to address the dual challenges of viral evolution and rapidly waning antibody levels.

ELI-005-induced T cell responses in mice displayed significant peak expansion post-boost in spleen, peripheral blood, and lung; sites important for early detection of viral infection both systemically and at the point of viral entry. Importantly, upon contraction, ELI-005-induced memory T cell responses were maintained in mice for >32 weeks and exhibited profound expansion upon recall. Long-lived anti-viral T cell responses have previously been observed in human subjects following natural infection and immunization including with the original SARS coronavirus, where T cell memory was present for years following infection^40^. Through early viral recognition, immediate execution of anti-viral effector function, and rapid expansion, these responses provide key contributions to long-term protection against viral progression, and onset of severe disease. Consistent with this, we observed that long-lived memory T cell responses in mice were rapidly expanded to peak levels upon re-exposure to antigen. Moreover, the presence of these persistent sentinel T cell populations in lungs and spleen may provide an effective source of viral surveillance in locations where respiratory viral entry or systemic signs of nascent infection could be readily detected.

In NHPs as in mice, ELI-005-induced T cells were numerous in peripheral blood confirming translation of ELI-005 activity to a model predictive of human immunity. Further, broad viral reactivity, with similar levels of cytokine-production were observed following stimulation with WH-01, Beta, Delta, and Omicron epitopes. This is consistent with epitope-mapping studies of human RBD-specific T cell responses showing that while viral variants commonly develop point mutations associated with partial evasion of antibody neutralization, most T cell-specific epitopes are maintained, representing a more consistent target for durable immune recognition^41^.

Effective accumulation of vaccine components in draining lymph nodes allows for potent coordination of innate immune mechanisms controlling the quality and potency of subsequent adaptive immunity. While many molecular adjuvants have proven immunomodulatory activity, their potency *in vivo* is often limited by poor exposure to the primary sites of immune surveillance and orchestration^19^. Instead, rapid, size-dependent clearance from the site of injection through the systemic circulation, and inefficient passive uptake into afferent lymphatics leads to biodistribution away from lymph nodes, toward immunologically irrelevant sites or tolerance promoting organs^19^. AMP-modification has proven to overcome this challenge by facilitating the physical association of CpG with tissue-resident albumin, a chaperone protein of optimal size (∼65 kDa) for clearance through lymphatic drainage rather than capillary absorption^21^. In practice, we observed that enhanced delivery of AMP-CpG to lymph nodes induced pleiotropic effects on the local innate immune response that correlated with significant improvements in downstream cellular and humoral immunity.

Compared to dose-matched vaccination with soluble CpG in mice, AMP-CpG triggered significant reprogramming of the lymph node transcriptome following administration. Previous reports have confirmed diminished or undetectable levels of soluble CpG in lymph nodes following immunization^21^. Here, soluble CpG induced a striking lack of effect on the lymph node transcriptome whereas AMP-CpG induced clear differential transcript levels marked by early and persistent chemokine upregulation, followed by enhancement of mechanisms underlying antigen presentation, pathogen recognition, anti-viral response, and APC engagement indicating a robust recruitment and coordination of innate immunity. The transcriptional signature observed in NHP lymph nodes exhibited similar patterns, suggesting shared mechanisms of activation in a model predictive of human activity, despite the more restricted expression pattern of TLR-9 in NHPs relative to mice. The observed signatures of transcriptional reprogramming were corroborated in mice by proteomic profiling where soluble CpG supported only transient increases in a limited group of chemokines. In contrast, a similar early increase in chemokine levels induced by AMP-CpG persisted alongside robust increases in nearly all analytes evaluated. Further mechanistic differentiation of soluble and AMP-CpG was observed through enumeration of activated and cytokine producing immune cell lineages where AMP-CpG induced costimulatory marker expression and cytokine production patterns similar to those seen in proteomic assessments, and indicative of a comprehensive coordination of innate immune cells to establish the lymph node inflammatory milieu. AMP-CpG induced significantly enhanced APC activity marked by DC and macrophage costimulation and cytokine production essential to optimal activation of T cells. Taken together, these observations provide evidence for the importance of AMP-modification to enhance CpG delivery to lymph nodes to realize the immunomodulatory potential of TLR-9 agonism to coordinate innate and adaptive immunity.

TLR-9 engagement by type B CpG DNA is predominantly thought to induce maturation of B cells and plasmacytoid DCs with lesser ability to promote production of Type I interferons relative to other CpG subclasses (reviewed in REF ^42^). However, in the case of AMP-CpG, the development of the lymph node inflammatory proteomic milieu was almost fully abrogated through blockade of IFNAR-1, suggesting a critical role for Type I interferons in the AMP-CpG-mediated coordination of acute inflammatory responses in the lymph nodes. While the mechanisms underlying this observation are not the subject of this study, it is notable that Type I interferon production by Type A CpG is dependent on macromolecular assembly of palindromic DNA sequences which drive signal transduction through TLR-9 clustering^43^. AMP-modification of CpG is known to promote insertion into lipid bilayers to densely decorate cell membranes^44,45^ in a pattern which might resemble the self-assembled structures critical to Type A CpG interferon responses. We speculate that decoration of the endosomal lumen-facing membrane leaflet by AMP-CpG could contribute to enhanced Type I interferon signaling resulting from TLR-9 clustering.

Despite the authorization, approval, and widespread uptake of effective vaccines throughout the developed world, global access has been low with challenges related to cold chain storage and distribution limiting availability in lower income countries^46^. Improvements in global vaccine coverage are thought to be an important step towards limiting further variant emergence resulting from community spread in populations with poor vaccine access. Such variants, including the recent example of Omicron, continue to prolong the pandemic, driving global waves of infection. Evaluation of ELI-005 following exposure to room temperature or refrigerated conditions demonstrated reduced temperature sensitivity with comparable levels of T and B cell-mediated immunogenicity maintained in the absence of continuous ultra-cold storage. The potential for reduced reliance on energy and logistically intensive transportation infrastructure for distribution of ELI-005 could allow for improved global vaccine access.

In summary, these studies provide insight into the potential of AMP-modification for directed lymph node delivery of CpG DNA to induce immunogenic responses through comprehensive programming of lymph node innate immunity. Application of this approach to subunit vaccine development offers a strategy to provide balanced, potent, and persistent cellular and humoral immunity against infectious disease antigens as exemplified by validation of ELI-005 immunogenicity in mice and NHPs. The potential for simple formulation development through admixture with protein subunit immunogens further suggests a potential for rapid application across a breadth of potential indications with high unmet medical need.

## Acknowledgements

We thank D. M. Lidgate for expert medical writing assistance, and C. S. Davis at CSD Biostatistics Inc., A. Bot at Capstan Therapeutics and D. J. Irvine at M.I.T. for helpful advice and discussion.

## Funding

We acknowledge support from Elicio Therapeutics.

## Author contributions

L.M.S., A.J., M.P.S., E.P., L.K.M. and P.C.D. drafted the manuscript. M.P.S., L.K.M., C.M.H. and P.C.D. provided supervision, and reviewed and finalized the manuscript draft. L.M.S., M.P.S., and P.C.D. designed the murine immunogenicity, and transcriptomics studies and analyzed the data. L.M.S., E.P., and M.P.S. conducted the murine immunogenicity, and transcriptomics studies. A.J. and P.C.D. designed the murine proteomics and ex vivo cell characterization studies and analyzed the data. A.J. and E.P. conducted the murine proteomics and ex vivo cell characterization studies. L.K.M, J.F., F.V., and P.C.D. designed the NHP studies. C.C. conducted the NHP sample collection. L.M.S., M.P.S., E.P. and L.K.M. conducted the NHP sample analyses. L.K.M., L.M.S., M.P.S., and P.C.D. analyzed the NHP data. All authors reviewed and approved the version for publication.

## Competing Interests

L.M.S., A.J., M.P.S., E.P., C.M.H., L.K.M, and P.C.D. are employees of Elicio Therapeutics and, as such, receive salary and benefits, including ownership of stock and stock options from the company. P.C.D., M.P.S., L.M.S., and C.M.H. have an Amphiphile SARS-CoV-2 vaccine patent pending to Elicio. The authors declare no other competing interests.

Correspondence and requests for materials should be addressed to P.C.D.

---

**Supplementary Fig. 1.**
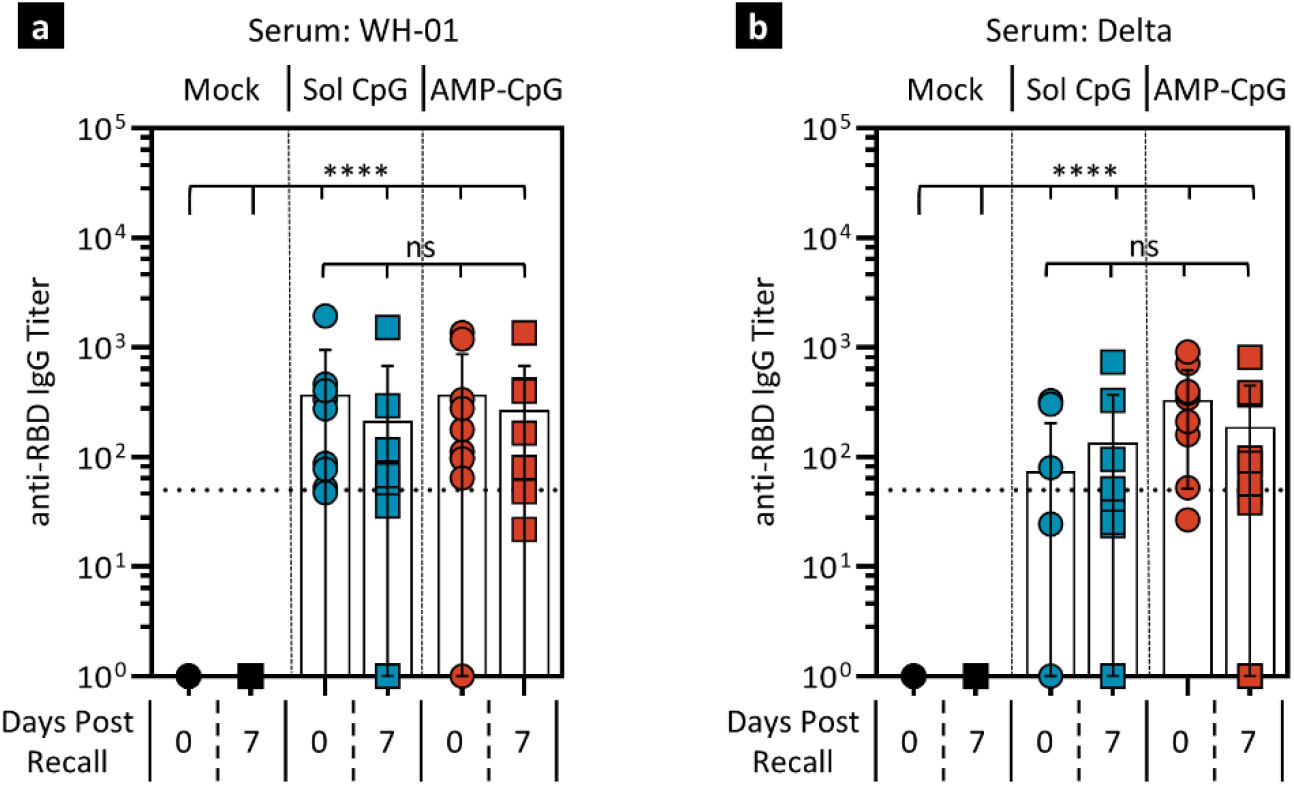
Serum antibody responses induced following SARS-CoV-2 Spike RBD immunization in mice. C57Bl/6J mice (n = 10) were immunized twice with 10 µg WH-01 RBD protein and 1 nmol soluble or AMP-CpG. 30 weeks post dose 2, mice were challenged subcutaneously with 10 µg WH-01 RBD protein. **a-b** Blood serum titers of anti-SARS-CoV-2 RBD antibodies were assayed 1 day before and 7 days after antigen challenge for WH-01 (**a**) and Delta (**b**). Values depicted are means ± standard deviation. *ns, not significant, ****p < 0*.*0001* by two-sided t-test applied to antibody titers.

**Supplementary Fig. 2.**
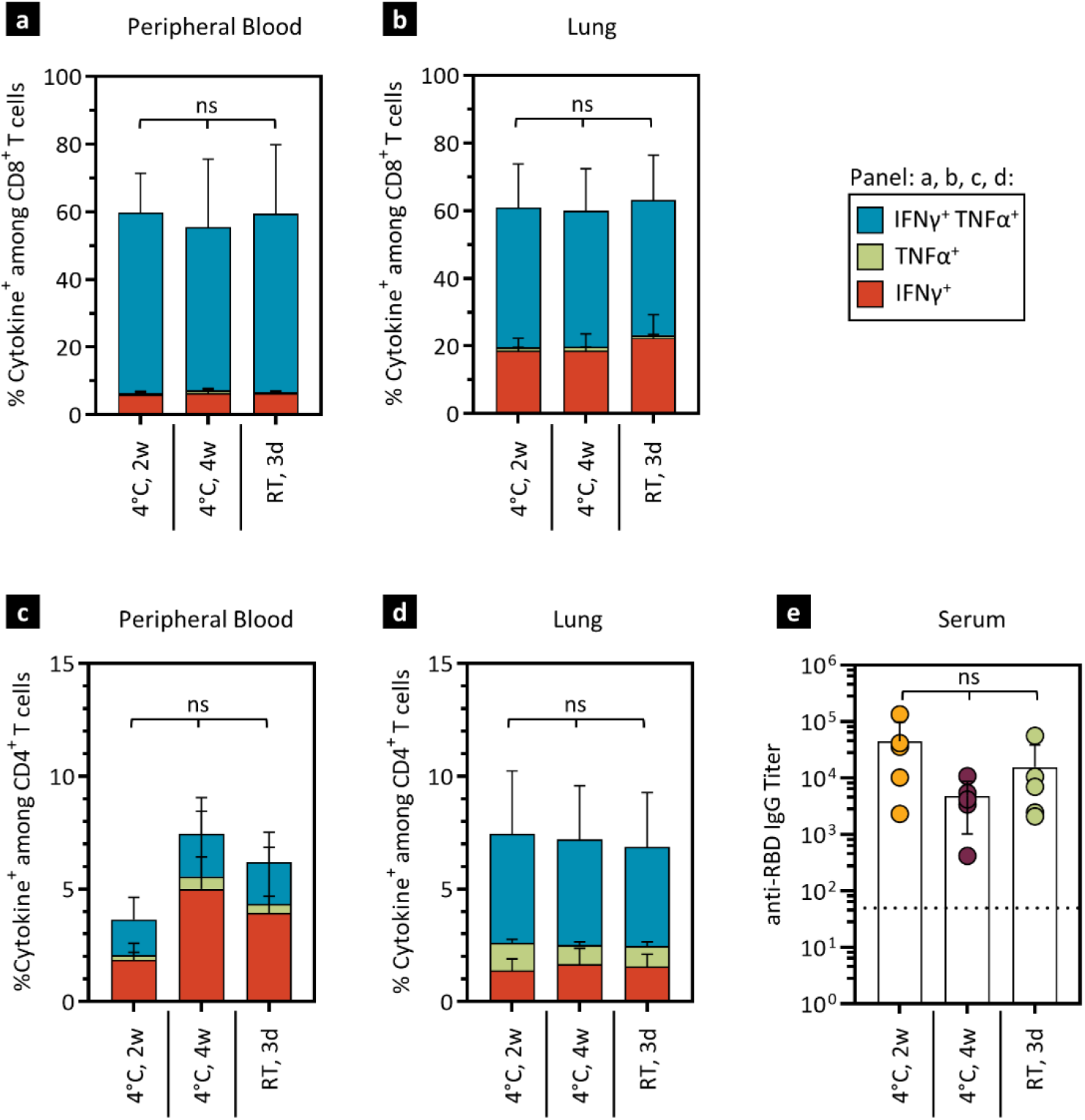
ELI-005 retains immunogenicity upon storage at refrigerated or ambient temperatures. ELI-005 vaccine doses, consisting of 10 µg WH-01 RBD protein and 1 nmol AMP-CpG, were admixed and stored at 4°C or 22°C (RT, room temperature) for the indicated period. C57Bl/6J mice (n=5) were immunized twice at week 0 and 2 and assayed 7 days after. CD8^+^ T cells from peripheral blood (**a**) and perfused lung (**b**), as well as CD4^+^ T cells (**c-d**) from those tissues were stimulated overnight with overlapping WH-01 RBD peptides and assayed for intracellular cytokines by flow cytometry. **e** 7 days post booster dose, blood serum was analyzed for anti-SARS-CoV-2 RBD antibody titers against WH-01 RBD. *ns, not significant*, by two-sided t-test applied to T cell frequencies and antibody titers.

**Supplementary Fig. 3.**
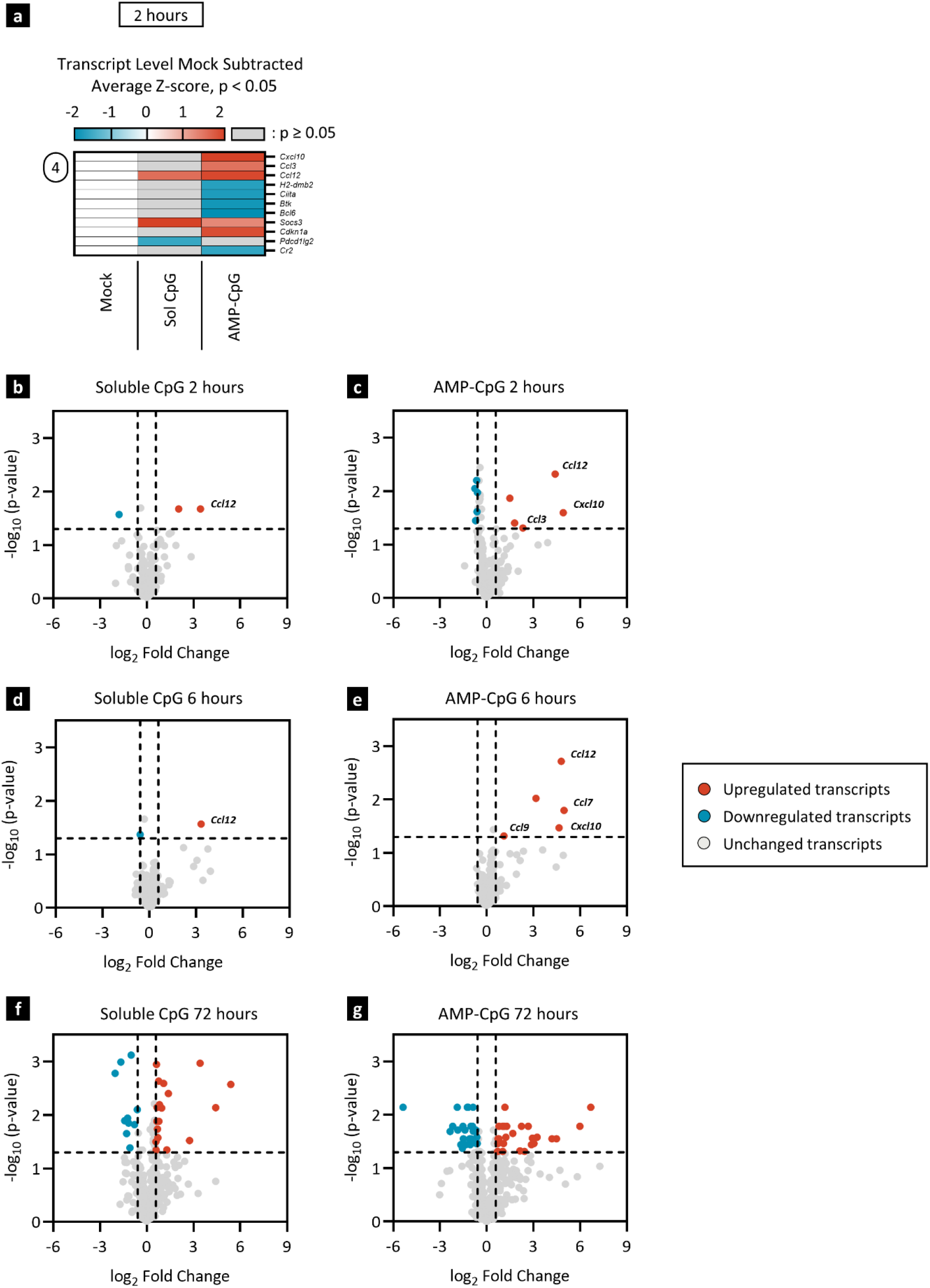
AMP-CpG immunization induces lymph node transcriptional reprogramming reflecting APC recruitment with increased potential for antigen processing and presentation. **a** Heatmap representation of whole lymph node mRNA analyzed by NanoString nCounter® Mouse Immunology Panel. Shown are mock-subtracted, average Z-scores of gene transcript levels significantly (p < 0.05) downregulated (≥ -1.5-fold change, blue) or upregulated (≥ 1.5-fold change, red) at 2 hours post injection relative to mock immunization. Insignificant values with p ≥ 0.05 are shown in gray. Gene groups follow the same numbering scheme as in **fig 3. b-g** Volcano plot representation of log-transformed soluble CpG and AMP-CpG transcript values at 2 hours (**b-c**), 6 hours (**d-e**), and 72 hours (**f-g**) post injection representing data from **supplementary figure 3a**, and figure **3b** and **3f**, respectively. Mock vaccines contained PBS vehicle only. Dotted horizontal line represents significance threshold of p = 0.05; vertical dotted lines represent fold-change limits of ± 1.5-fold change. Statistical analysis was performed using Rosalind software.

**Supplementary Figure 4.**
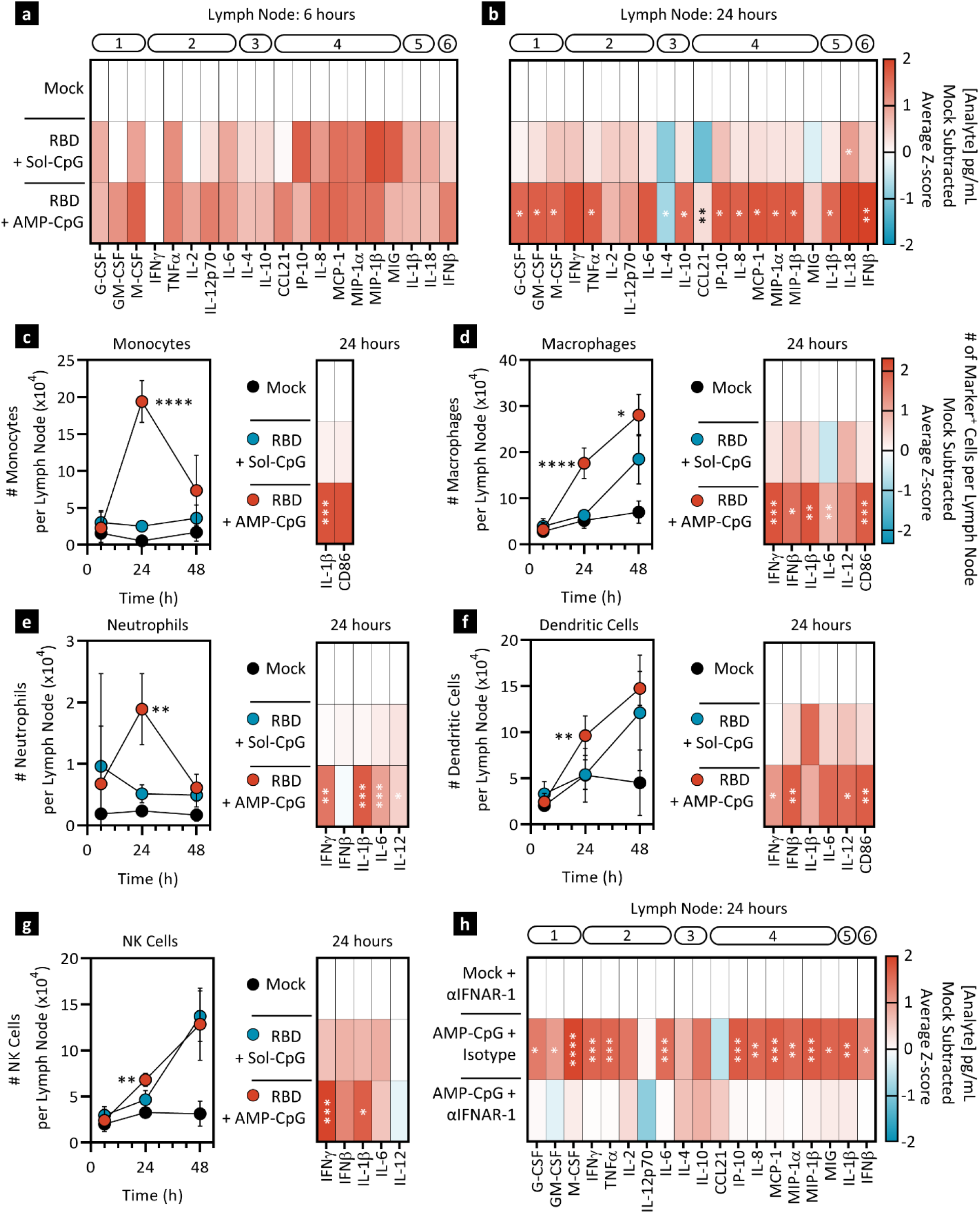
Additional statistical analysis for fig 4. Data reported in **fig 4** were re-analyzed to determine statistical significance between groups immunized with soluble and AMP-CpG, whereas statistical analysis in fig 4 compared to mock treatment. Values depicted are mean ± standard deviation. *\*p < 0*.*05; **p < 0*.*001; ***p < 0*.*0005; ****p < 0*.*0001* by non-parametric two-tailed T test comparing immunization with soluble CpG to AMP-CpG.

**Supplementary Fig. 5.**
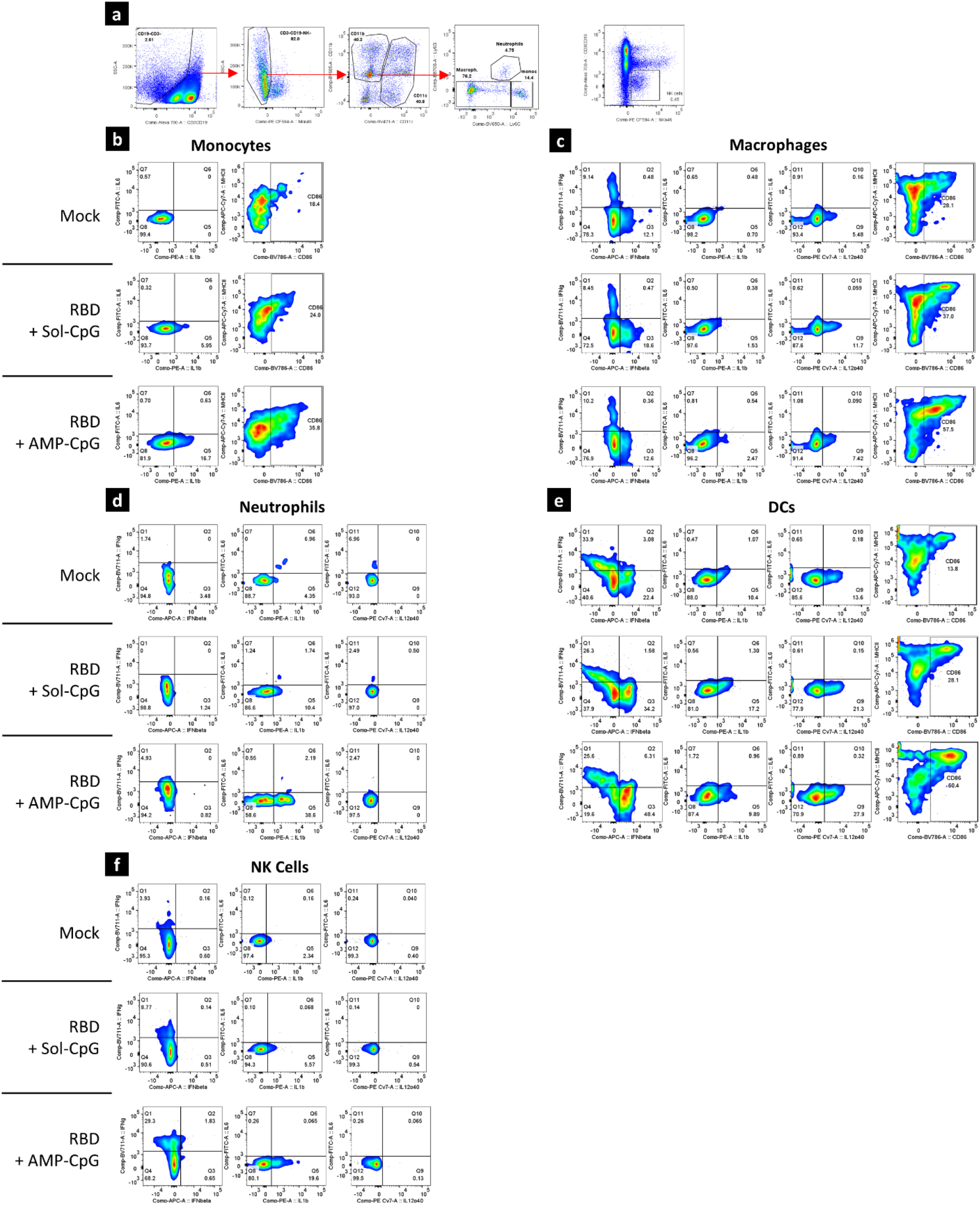
Supplementary data for fig 4. Flow cytometric analysis strategy and example scatter plots for surface marker and intra-cellular cytokine expression induced in innate immune cell lineages.

**Supplementary Fig. 6.**
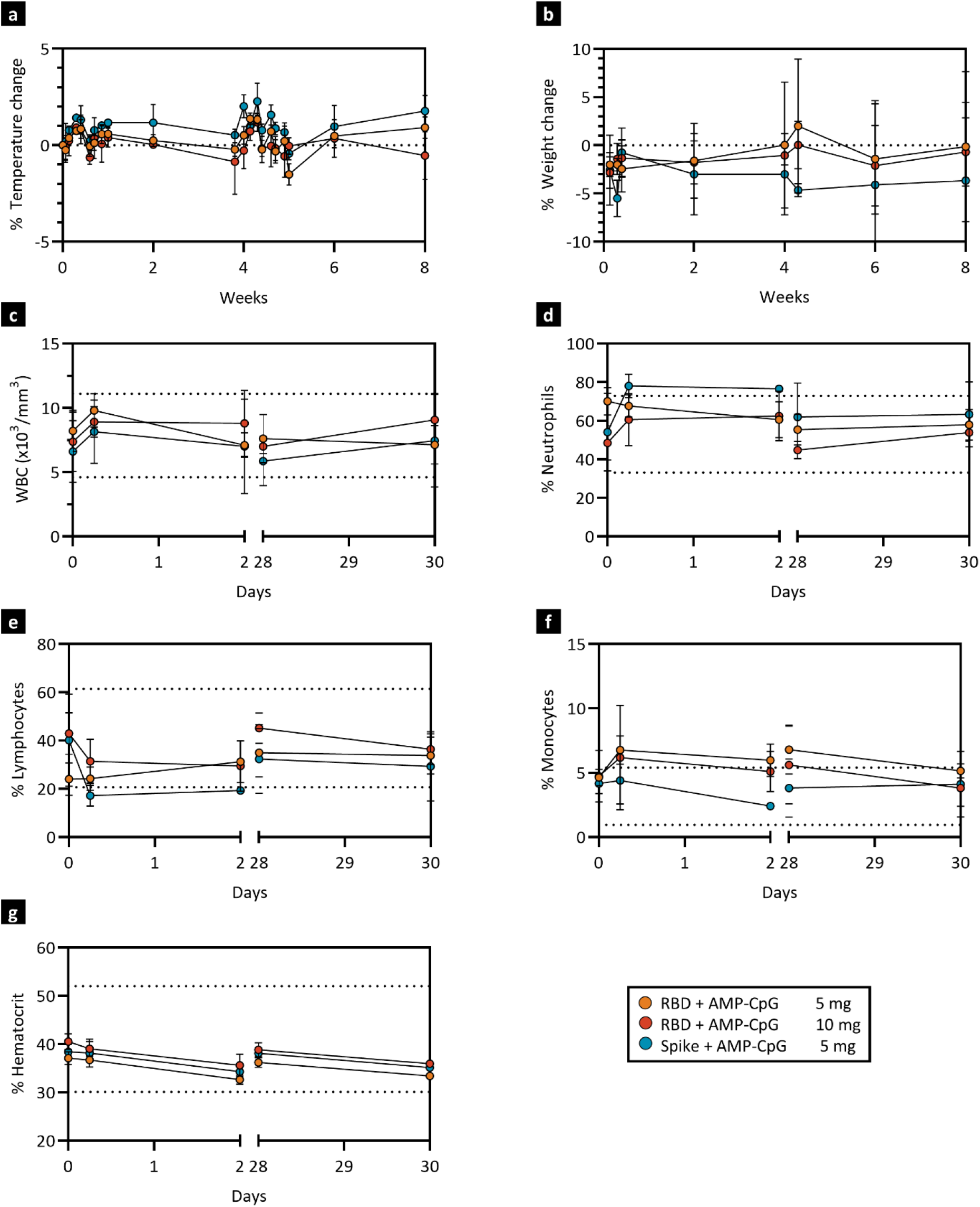
ELI-005 vaccination is safe and non-toxic in NHP. Rhesus macaques (n=2/3) were immunized at week 0 and 4 with 140 µg WH-01 RBD protein admixed with either 5 mg or 10 mg of AMP-CpG, or 140 µg full WH-01 Spike protein admixed with AMP-CpG at 5 mg. Vital signs were recorded. Shown are percent change for temperature (**a**) and weight change (**b**) from baseline. Sera was collected at multiple timepoints for assessment in a hematology complete blood count panel. Shown are white blood cells (WBC) (**c**), % neutrophils (**d**), % lymphocytes (**e**), % monocytes (**f**), and % hematocrit (**g**). Dotted lines indicate reference value ranges.

## METHODS

### Vaccine components

For mouse studies, vaccines consisted of 10 µg SARS-CoV-2 Spike RBD WH-01 protein (GenScript, cat# Z03483), combined with 1 nmol AMP-CpG-7909 (AMP-CpG, Avecia), or soluble CpG 7909 (soluble CpG, InvivoGen, tlrl-2006,). Mock treatment groups received a matching dose of adjuvant in the absence of antigen or were treated with vehicle alone (phosphate-buffered saline, PBS), as indicated. Antigen rechallenge doses consisted of 10 µg SARS-CoV-2 Spike RBD WH-01 protein, administered subcutaneously bilaterally into the tail base. For experiments testing storage temperature conditions, doses were admixed and stored at 4°C or room temperature (22°C) for the indicated time. For NHP studies, vaccines consisted of either SARS-CoV-2 Spike RBD WH-01 protein (GenScript; cat# Z03483) or SARS-CoV-2 WH-01 Spike protein (Acro Biosystems, SPN-C5H9) at 140 µg admixed with AMP-CpG at either 5 or 10 mg per dose.

### Reagents

For stimulation of *ex vivo* cell samples, OLP peptide pools of 15mers with 11 amino acid overlap were generated spanning the SARS-CoV-2 Spike RBD (R319-S591, GenScript). Sequences that contained VOC mutations were exchangeable with the corresponding mutated peptides due to a modular OLP pool design.

### Animals

All animal studies were carried out under an institute-approved Institutional Animal Care and Use Committee (IACUC) protocol following federal, state, and local guidelines for the care and use of animals. For mouse studies, female 6- to 8-week-old C57BL/6J mice were purchased from the Jackson Laboratory (Bar Harbor, ME). The indicated antigen-adjuvant combinations were administered into mice subcutaneously at the base of the tail (bilaterally, 50 µl each) at weeks 0 and 2. Peripheral blood samples were collected as indicated 7 days post second dose, and 1 day prior to and 7 days post antigen challenge where applicable. Spleen and lung tissue was collected either 7 days post second dose or 7 days post antigen challenge. Lungs were harvested following perfusion with 10 mL of PBS into the right ventricle of the heart. Lung tissue was physically dissociated and digested with RMPI 1640 media containing collagenase D (1 mg/mL) and deoxyribonuclease I (25 U/mL).

For NHP studies, 8 outbred, Indian-origin, 4-5 year old female rhesus macaques (*Macaca mulatta*) were randomly allocated into 3 groups of 2 or 3 animals. All animals were housed at New Iberia Research Center (New Iberia, LA). Animals received immunizations subcutaneously into the upper thigh at week 0 and 4. Blood samples were obtained at baseline and throughout the study and processed for PBMCs. Serum was collected at baseline, 0, 2, 6, 24, and 48 hours, and 2, 4, 6, and 8 weeks post priming dose. Lymph node FNAs were collected from the inguinal lymph nodes at weeks 0 and 4 for flow cytometric analysis of lymph node resident cells and 24 hours post week 10 boost for NanoString analysis.

### Cell activation and cytokine determination by ICS

For mouse vaccine immunogenicity studies, ICS analysis was performed as described previously (Steinbuck *et al*., 2021).

For mouse innate immune response analyses, surface activation marker staining and ICS analysis was performed on fixed/permeabilized (BD, cat# 554714) single cell suspensions of inguinal lymph nodes collected 6-48 hours post immunization. Live/Dead fixable stain (Aqua, Invitrogen, cat# L34966) was used to exclude dead cells. Cells were stained with: CD11b (BV605, clone: M1/70, BioLegend), CD11c (BV421, clone: N418, BioLegend), CD3 (AF700, clone: 17A2, BioLegend), CD19 (AF700, clone: 6D5, BioLegend), Ly6C (BV650, clone: HK1.4, BioLegend), Ly6G (PcP-Cy5.5, clone: 1A8, BioLegend), NKp46 (PE-Dazzle594, clone: 29A1.4, BioLegend), MHCII (APC-Cy7, clone: M5/114.15.2, BioLegend), CD86 (BV785, clone: GL-1, BioLegend), IFNγ (BV711, clone: XMG1.2, BioLegend), IL12p40 (PECy7, clone: C15.6, BioLegend), IL1β (PE, clone: NJTEN3, Invitrogen), IL6 (AF488, clone: MP5-20F3, Invitrogen), and IFNβ (APC, ASSAYPRO, cat# 32183-05161T). Sample acquisition was performed on BD FACS Symphony and data were analyzed with BD FlowJo V10 software.

Macrophages were defined as CD3^-^/CD19^-^/NKp46^-^/CD11c^-^/Ly6G^-^/CD11b^+^/Ly6C^low^; monocytes were defined as CD3^-^/CD19^-^/NKp46^-^/CD11c^-^/Ly6G^-^/CD11b^+^/Ly6C^high^; DC were defined as CD3^-^/CD19^-^/NKp46^-^/CD11b^-^/CD11c^+^; Neutrophils were defined as CD3^-^/CD19^-^/NKp46^-^/CD11c^-^/CD11b^+^ /Ly6C^med^/Ly6G^+^; NK cells were defined as CD3^-^/NKp46^+^. Data was expressed as total cell number per lymph node over time, or as Z-scores of the number of cells that were positive for the respective cytokine or surface marker.

For NHP studies, frozen PBMCs were thawed and rested overnight. 10^6^ PBMCs/well were resuspended in R10 media supplemented with anti-CD49d monoclonal antibody (clone: 9F10, BD), anti-CD28 monoclonal antibody (clone: CD28.2, BD), and Golgi inhibitors monensin (Fisher Scientific, cat# NC0176671) and brefeldin A (Fisher Scientific, cat# 50-112-9757) and incubated at 37°C for 8 hours, then maintained at 4°C overnight. The next day, cells were surface-stained with antibodies against CD4 (PE-Cy5.5, clone: S3.5, Invitrogen), CD8 (AF647, clone: RPA-T8, BioLegend), CD45RA (FITC, clone: 5H9, BD), CCR7 (BV650, clone: G043H7, BioLegend), and aqua live/dead dye (Invitrogen, L34957), and subsequently fixed with BD CytoFix/CytoPerm (BD, 554714). Cells were further stained with antibodies against CD3 (APC-Cy7, clone: SP34-2, BD), CD69 (ECD, clone: TP1.55.3, Beckman Coulter), IFNγ (AF700, clone: B27, BioLegend), IL-2 (BV421, clone: MQ1-17H12, BioLegend), IL-4 (PE, clone: 8D4-8, BioLegend), TNFα (BV605, clone: MAb11, BioLegend), and IL-17A (PE-Cy7, clone: BL168, BioLegend). Cells fixed in 1.5% formaldehyde were acquired on a BD FACS Symphony and data were analyzed with BD FlowJo V10 software.

### Mouse antigen-specific tetramer staining of peripheral blood cells

MHC-tetramer staining on mouse samples was performed using an RBD-PE tetramer specific for sequence VNFNFNGL (NIH Tetramer Core Facility at Emory University, cat# 54971), and antibodies against CD8a (APC, clone: 53-6.7, eBioscience), CD3 (APC-Cy7, clone: 17A2, BD), CD44 (PE-Cy7, clone: IM7, eBioscience), and CD62L (FITC, clone: MEL-14, eBioscience). A Live/Dead fixable (aqua) dead cell stain kit (Invitrogen, cat# L34966) was used to evaluate viability of the cells during flow cytometry. Cells were permeabilized and fixed with a transcription factor staining buffer set (eBioscience, cat# 00-5523-00). Sample acquisition was performed on a BD FACSCanto II and data were analyzed with BD FlowJo V10 software.

### Enzyme linked immunospot (ELISpot) Assay

For mouse studies, spleens were collected and processed into single-cell suspensions. Red blood cells were lysed in ACK lysis buffer (Quality Biological Inc., cat# no. 118156101). ELISpot assays for IFNγ were performed as instructed by the manufacturer using the Mouse IFNγ ELISpot Set (MabTech, cat# Q00824-1×20210623DS). 0.1×10^6^ mouse splenocytes were plated into each well and stimulated overnight with 0.2 µg per peptide per well of RBD-derived overlapping peptides (GenScript).

For NHP studies, ELISpot assays were performed using Monkey IFNγ ELISpotPLUS kits (MabTech, cat# 3241M-4HPW). Precoated 96-well ELISpot plates were blocked with RPMI + 10% FBS for 2 hours at room temperature. 0.2×10^6^ PBMCs were plated into each well and stimulated overnight with 0.4 ug per peptide per well of RBD-derived OLPs for WH-01, Beta and Delta variants. The spots were developed based on the manufacturer’s instructions. In both animal models, PMA (50 ng/mL) and ionomycin (1 µM) were used as positive controls, and RPMI + 10% FBS with DMSO was used as the negative control. Spots were scanned and quantified using an S6 ImmunoSpot analyzer (CTL).

### Enzyme linked immunosorbent assay (ELISA) for antibody titers

For mouse studies, ELISA assays were performed as previously described (Steinbuck *et al*., 2021). For NHP assays, ELISA plates were coated with 100 ng/well of SARS-CoV-2 RBD antigen. The detection antibody used was horseradish peroxidase (HRP)–conjugated goat anti-human IgG (H+L) (ThermoFisher, cat# SA5-10283) at a 1:2000 dilution. The WHO International Standard for anti-SARS-CoV-2 immunoglobulin (anti-RBD IgG High 20/150, NIBSC) was used for reference values. Serum titers were determined at an absorbance cutoff of 0.5 OD and converted into binding antibody units/mL (BAU/mL) using the WHO standard.

### Gene transcript analysis by Nanostring

For mouse studies, inguinal lymph nodes were harvested from immunized C57BL/6J mice at the indicated time points, processed into single cell suspensions, and lysed with RLT buffer (Qiagen, cat# 79216). Transcriptional profiles of immune signaling were generated using the nCounter Mouse Immunology Panel of 568 mouse immune response genes (NanoString Technologies). For NHP studies, lymph node FNA were assessed using the nCounter NHP Immunology Panel of 770 macaque immune response genes (NanoString Technologies). Transcriptional responses were assessed with nSolver software v4.0 (NanoString Technologies) and differential gene expression was carried out using ROSALIND software.

### Mouse lymph node proteomics

To determine the cytokine/chemokine content of lymph nodes, animals were vaccinated and inguinal lymph nodes were collected 6-48 hours post immunization. For IFNAR-1 blockade, an antagonistic mAb (clone: MAR1-5A3, BioXcell) or an isotype control (clone: MOPC-21, BioXcell) were injected intraperitoneally 24 hours prior to immunization. Protein Extraction Buffer (Invitrogen, cat# EPX-9999-000) containing Mini protease inhibitor cocktail (Roche, cat# 53945000) and HALT phosphatase inhibitors (Thermo Fisher, cat# 78442) was added to the intact lymph nodes prior to homogenization using a TissueLyser II (Qiagen). Centrifugation-cleared lysates were analyzed with Luminex Cytokine and Chemokine kits (EMDMillipore, cat# MCYTOMAG-70K and MECY2MAG-73K).

### NHP antigen-specific GC B cell analysis in lymph node FNAs

Biotinylated RBD protein (Acro Biosystems, cat# SPD-C82E9) was complexed with fluorochrome-conjugated streptavidin APC (BioLegend, cat# 405207). Lymph node cells from FNA were incubated with the RBD-fluorochrome complex and subsequently stained with aqua live/dead dye (Invitrogen, L34957), anti-human IgM (FITC, clone: G20-127, BD), anti-human IgG (PE-Cy7, clone: G18-145, BD), anti-human CD3 (AF700, clone: SP34-2, BD), anti-human PD-1 (BV650, clone: EH12.1, BD), anti-human CD20 (PE/Dazzle 594, clone: 2H7, BioLegend), anti-human CD4 (APC-Cy7, clone: OKT4, BioLegend) and anti-human CXCR5 (PcP-eF710, clone: MU5UBEE, ThermoFisher). Cells were fixed/permeabilized using Transcription Factor Staining Buffer Set (ThermoFisher, cat# 00-5521-00) and further stained with anti-human Bcl-6 (PE, clone: 7D1, BioLegend) and anti-human Ki-67 (BV421, clone: 11F6, BioLegend). Sample acquisition was performed on a BD FACS Symphony and data were analyzed with BD FlowJo V10 software.

### SARS-CoV-2 pseudovirus neutralization assay for NHP sera

The SARS-CoV-2 pseudovirus assay was performed by Genecopoeia as previously described^47^. SARS-CoV-2 Spike-pseudotype lentiviruses from the Washington 1 (WA1) D614G, Delta and Omicron variants were used. Briefly, a HEK293T cell line overexpressing ACE2 and TMPRSS2 was seeded at a density of 1.2×10^4^ cells/well overnight. 3-fold serial dilutions of heat inactivated serum samples were prepared and mixed with 50 µL of pseudovirus. The mixture was incubated at 37°C for 1 hour before adding to the HEK-293T-hACE2 cells. After 72 hours, the cells were lysed, and the firefly luciferase activity was determined. SARS-CoV-2 neutralization titers were defined as the sample dilution at which a 50% reduction in RLU was observed relative to the average of the virus control wells. Convalescent serum samples and plasma samples from patients who had recovered from SARS-CoV-2 infection (COVID-19) were obtained from US Biolab (Rockville, MD) and ALLCELLS (Alameda, CA). All samples were received and stored frozen at −80°C until analysis.

### Statistics

For comparing two experimental groups, two-tailed t-test analysis was utilized when normal distribution and homogeneity of variance, determined by Levene’s Test, were established. Where these assumptions did not apply, the Mann-Whitney test was used instead. For comparison of multiple groups, ordinary one-way ANOVA was used to compare experimental groups. NanoString statistical analysis was performed using Rosalind software.

